# Aberrant MNX1 Expression Associated with t(7;12)(q36;p13) Pediatric Acute Myeloid Leukemia Induces the Disease Through Altering Histone Methylation

**DOI:** 10.1101/2022.06.10.495634

**Authors:** Ahmed Waraky, Anders Östlund, Tina Nilsson, Dieter Weichenhan, Pavlo Lutsik, Marion Bähr, Joschka Hey, Jenni Adamsson, Mohammad Hamdy Abdelrazak Morsy, Susann Li, Linda Fogelstrand, Christoph Plass, Lars Palmqvist

**Author notes:** Corresponding author Lars Palmqvist MD, PhD, Institute of Biomedicine, University of Gothenburg Sahlgrenska University Hospital, SE-413 45 Gothenburg, Sweden. **Authors contribution** LP, AÖ and AW designed the research study. AW, AÖ, TN, DW, PL, JH, JA, SL, MHM performed laboratory work and result analysis. AW, LF, CP and LP analyzed the combined data and wrote the paper.

## Abstract

Certain subtypes of acute myeloid leukemia (AML) in children have inferior outcome. One of these has a translocation t(7;12)(q36;p13) leading to a *MNX1*::*ETV6* fusion along with high expression of MNX1. Here we identified the transforming event in this AML and possible ways of treatment. Only MNX1 was able to induce AML in mice, with similar gene expression and pathway enrichment to t(7;12) AML patient data. Importantly, this leukemia was only induced in immune incompetent mice using fetal but not adult hematopoietic stem and progenitor cells. The restriction in transforming capacity to cells from fetal liver is in alignment with t(7;12)(q36;p13) AML being mostly seen in infants. Expression of MNX1 led to increased histone 3 lysine 4 mono-, di- and trimethylation, reduction in H3K27me3, accompanied with changes in genome-wide chromatin accessibility and genome expression, likely mediated through MNX1 interaction with the methionine cycle and methyltransferases. MNX1 expression increased DNA damage, depletion of the Lin-/Sca1+/c-Kit+ population and skewing toward the myeloid lineage. These effects, together with leukemia development, was prevented by pretreatment with the S-adenosylmethionine analog Sinefungin. In conclusion, we have shown the importance of MNX1 in development of AML with t(7;12), supporting a rationale for targeting MNX1 and downstream pathways.

## INTRODUCTION

Non-random cytogenetic aberrations are often involved in the development of acute myeloid leukemia (AML) and several aberrations can serve as diagnostic markers, prognosis predictors, and impact the choice of therapy ^1^. In AML diagnosed in children under the age of 24 months, a chromosomal translocation t(7;12)(q36;p13) with poor prognosis has been reported ^2^. There have been contradictory results on the incidence of t(7;12) but recent studies suggest the frequency in children <24 months to be 5-7% ^3, 4^. Similarly, different results have been reported regarding the prognosis where recent studies show 20%-43% 3-year event-free survival but with a high relapse rate, ranging from 57-80% ^3, 4^. However, the understanding of the mechanisms behind the leukemia transformation of t(7;12) AML remains poorly understood.

The chromosomal break points in t(7;12) have consistently been found to be located close to the Motor neuron and pancreas homeobox 1 (*MNX1)* gene on chromosome 7, and in introns 1 or 2 in the *ETV6* gene in chromosome 12 ^5^. The translocation leads to *MNX1* gene activation and in most reported cases also to a *MNX1::ETV6* fusion transcript consisting of exon1 of *MNX1* transcript variant 1 spliced to the remaining ETV6 exons, depending on the location of the break point in ETV6 ^6^.

*MNX1*, also known as Homeobox HB9 (*HLXB9*), belongs to the homeobox domain family of transcription factors, with previous studies showing the importance of *MNX1* in motor neuron development ^7^, pancreas development ^8, 9^and in hereditary sacral agenesis ^10^. *ETV6*, also known as *TEL* belongs to the ETS-family transcription factors. *ETV6* encodes a transcriptional repressor that plays a critical role in embryonic development and hematopoiesis, where it is essential for normal HSC function and generation of thrombocytes by megakaryocytes ^11^.

Translocations involving the chromosomal region of 12p13 that result in the rearrangements of the *ETV6* gene are one of the most observed chromosomal abnormalities in human leukemia with more than 30 reported translocations. These chromosomal translocations can induce leukemias through the ectopic expression of a proto-oncogene in the vicinity of a chromosomal translocation ^12^ or the constitutive activation of the partner protein ^13^. In addition, the formation of ETV6 fusion proteins can result in the modification of the original functions of the transcription factor ^14^, or loss of function of the fusion gene, affecting *ETV6* and the partner gene ^15^.

The role of the MNX1::ETV6 fusion protein in the development of AML with t(7;12) has not been established. It is also unclear whether the driver of leukemogenesis is the MNX1::ETV6 fusion protein or overexpression of MNX1. The aim of this study was to assess the transformation capacity and the molecular mechanism of the MNX1::ETV6 fusion and the ectopic expression of MNX1 *in vitro* and *in vivo* using murine transplantation models.

## MATERIAL AND METHODS (Complete materials and methods in supplement)

### Plasmid constructions

The *MNX1, ETV6* and MNX1::ETV6 fusion sequences are listed in Supplement table 1. An HA-tag (36bp) is introduced at the 5’ end and via a linker sequence of 24 bp attached to the separate gene sequences where the first ATG is removed. These constructs were cloned into the MSCV-IRES-GFP and MSCV-IRES-YFP vectors (Takara, Japan) #634401 under control of the viral LTR promoter.

### Generation of transduced murine bone marrow (BM) cells and transplantation assays

Mice were bred and maintained at the Gothenburg university laboratory for experimental biomedicine Animal Facility (Gothenburg, Sweden) in an SPF environment. Establishment and characterization of BM cell lines following transduction of BM cells with *MNX1, MNX1::ETV6, ETV6 or* empty vector Ctrl have been done as described previously ^16^. In brief, BM cell lines were established from BM cells previously treated with 5-fluorouracil (5-FU) from 8-12 weeks old (C57Bl/6) mice (Charles River Laboratories, Inc., Wilmington, Massachusetts, USA) for three days for adult bone marrow (ABM) or from fetal liver at embryonic days 14.5 (E14.5) and maintained in liquid culture (Dulbecco’s modified Eagle’s medium supplemented with 18% fetal bovine serum, 10 ng/mL human interleukin [IL]-6, 6 ng/mL murine IL-3, and 50 ng/mL murine stem cell factor). All culture media were obtained from Sigma and recombinant growth factors from Peprotech. To generate *MNX1, MNX1::ETV6, ETV6 or* empty vector Ctrl BM cell lines, the BM cells were transduced by cocultivation on irradiated (4000 cGy) E86 producers (ATCC, Manassas, VA, USA) for a period of 2 days in the presence of 5 μg/mL protamine sulfate (Sigma). Cells were sorted for GFP^+^ and/or YFP^+^ expression by flow cytometry analysis (fluorescence-activated cell sorting [FACS]) FACSAria (BD Biosciences) and maintained in culture for 5-7 days post transduction before transplantation in mice. Lethally (8.5 Gy) or sublethally (5.5 Gy) radiated 8-12 weeks old C57Bl/6 mice received the equivalent of 0.6×10^6^ rescue BM cells and/or 0.8-1×10^6^ transduced cells via tail vein injection. For immunocompromised NOD.Cg-*Kit*^*W-41J*^ *Tyr* ^+^ *Prkdc*^*scid*^ *Il2rg*^*tm1Wjl*^/ThomJ (*NBSGW*) *mice (*The Jackson Laboratory, Bar Harbor, ME, USA), cells were transplanted either with no radiation, or after lethal dose of (1.6 Gy) and sublethal dose of (0.9 Gy). Donor-derived engraftment and reconstitution were monitored by flow cytometry analysis for GFP^+^ and/or YFP^+^ expression in the peripheral blood of the transplants every 2 weeks. Mice were sacrificed using Isoflurane (Baxter, Deerfield, IL, USA). Blood counts were analyzed on a Sysmex KX-21 Hematology Analyzer (Sysmex, Norderstedt, Germany).

### Statistical analysis

Two-sided Student’s t-test were used for comparisons between different groups in all experiments, unless stated otherwise. Log-rank test was used to compare survival between mice groups. Mann-Whitney two-tailed U-test was used for comparison between t(7;12) patients and normal human bone marrow from Target cohort.

#### Ethics Statement

All animal experiments have been accepted by the Jordbruksverket and the animal ethics committee in Gothenburg, Dnr 5.8.18-17008/2021.

## RESULTS

### *MNX1* induces acute myeloid leukemia in hematopoietic cells of fetal origin

To investigate the leukemogenic potential of t(7;12), we transduced primary murine (C57BL/6) hematopoietic stem and progenitor cells (HSPC) from either adult bone marrow after 5-FU stimulation (ABM-HSPC) or fetal liver cells at E14.5 (FL-HSPC) with retroviral vectors for expression of the *MNX1*::*ETV6* fusion, *MNX1, ETV6* or empty vector (Ctrl). Expression of *MNX1, ETV6* and *MNX1*::*ETV6* was confirmed in both FL-HSPC and ABM-HSPC (Supplement figure 1a-c). The transduced cells were transplanted into lethally irradiated C57BL/6 mice with rescue bone marrow. Neither *MNX1* overexpression nor *MNX1::ETV6* in ABM-HSPC or FL-HSCP were able to induce leukemia in these mice, as assessed by survival, WBC, hemoglobin, blood smears and spleen size (Supplement figure 2a-d). However, when increasing the percentage of MNX1 chimerism through sublethal radiation with no rescue BM 20% of the mice exhibited signs of malignant transformation after transplantation with FL-HSPC, including high WBCs count, severe anemia and enlarged spleen (Supplement figure 2e-f & 3a-b). To further examine the leukemogenic potential of *MNX1* overexpression and the *MNX1*::*ETV6* fusion, we repeated the experiment using immune-compromised non-irradiated NOD.Cg-*Kit*^*W-41J*^ *Tyr* ^+^ *Prkdc*^*scid*^ *Il2rg*^*tm1Wjl*^/ThomJ (NBSGW) mice. Within 12-18 weeks after transplant, the *MNX1* mice showed clear signs of leukemia including pallor, weight loss, severe anemia, leukocytosis with a high percentage of *MNX1*-transduced cells, elevated blast cells in blood and bone marrow, enlarged spleen and liver infiltrated by leukemia (Figure 1a-d). Cells from bone marrow showed predominant expression of the c-Kit protein, but not the stem cell marker SCA-1 or more differentiated myeloid markers such as Mac-1, Ly6G1 and Ly6C1, suggesting a poorly differentiated myeloid leukemia (Figure 1d; Supplement figure 3d and 4a). To rule out that the leukemia development in immunocompromised mice was not due to enhanced transplantation efficiency, we reduced the chimerism of *MNX1* cells in NBSGW mice through the usage of lethal radiation and rescue bone marrow. These mice also developed leukemia but with a slightly longer latency than mice that were sublethally irradiated without rescue BM (Supplement figure 3d). Acute leukemia induction by MNX1 was confirmed by leukemia development after secondary transplant of BM from mice with primary leukemia, both in non-irradiated mice and in sublethally irradiated mice receiving rescue BM (Supplement figure 5a). When NBSGW mice were transplanted with ABM-HSPC transduced with *MNX1*, there were no signs of leukemia six months after the transplant (Figure 1e; Supplement figure 5c-d). Taken together, leukemogenesis was achieved by MNX1 only in the immunocompromised setting with fetal origin of leukemic cells.

**Figure 1.**
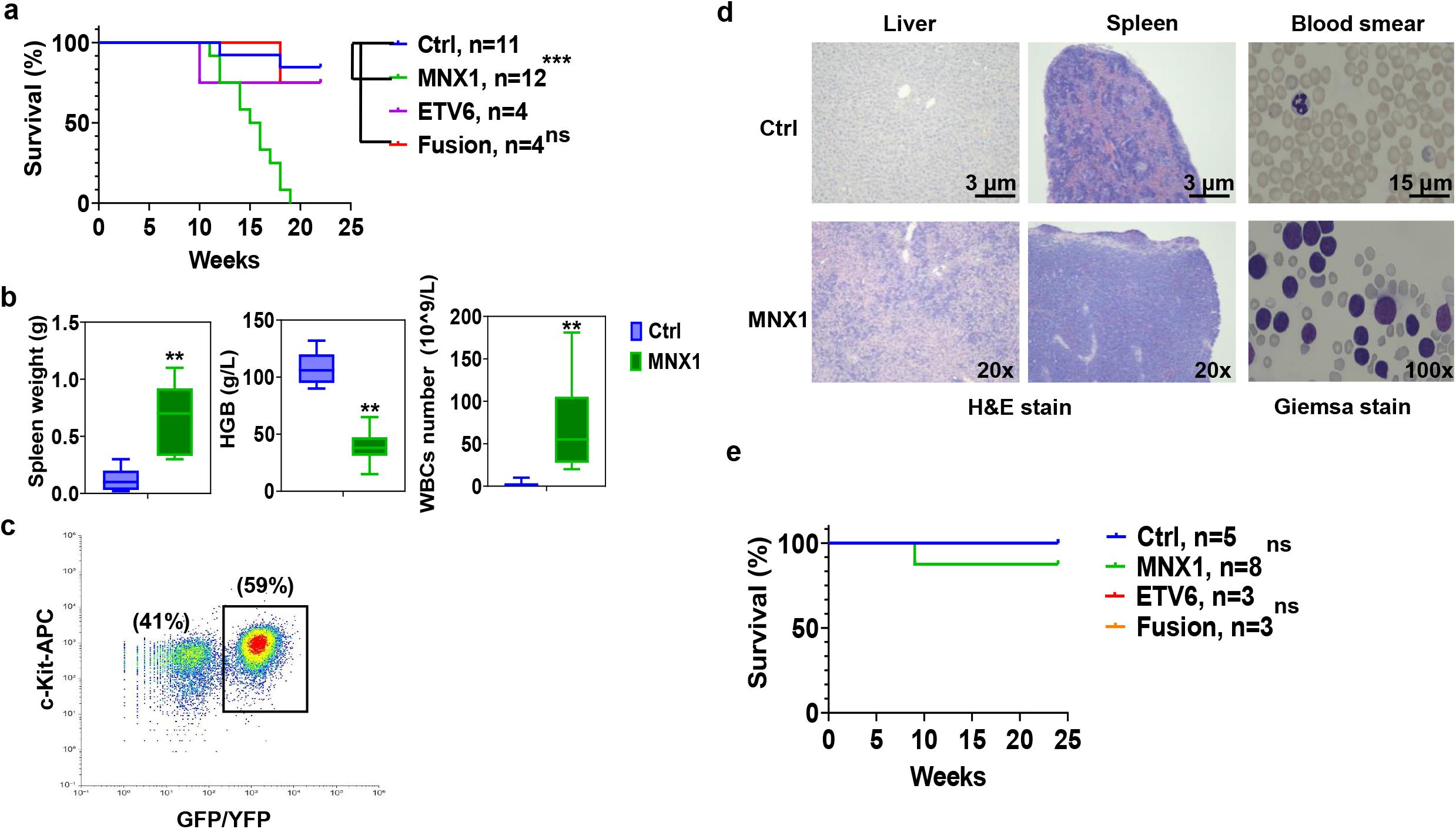
MNX1 induces AML in immunocompromised mice. Kaplan-Meier survival curve of (NBSGW) mice transplanted with FL-HSPC after retrovirus transduction with either ectopic expression of *MNX1* (green), *ETV6* (purple), *MNX1::ETV6* fusion (red) or empty vector control (blue). Results from the transplanted mice (control n = 11, *MNX1*: n = 12, *ETV6* and *MNX1::ETV6* n = 4) were analyzed using the log-rank test (***: significant at *P* ≤ 0.01, ns: non-significant) (a). Mice were euthanized when showing sign of disease or at the end of the experiment and analyzed for spleen weight, white blood cell number and hemoglobin concentration (b). Quantification of flow cytometry analysis of BM cells from (NBSGW) mice with c-Kit expression showing all events. GFP/YFP were used as indicative for MNX1 expression (c). Representative images of H- and E-stained formaldehyde fixed spleen and liver sections from control and MNX1 mice (d; left-panels). Representative images of Giemsa-stained peripheral blood smears from Ctrl and MNX1 mice (d; right-panels) (d). Kaplan-Meier survival curves of (NBSGW) mice transplanted with ABM cells with either ectopic expression of *MNX1, ETV6, MNX1::ETV6* fusion or empty vector control after sub-lethal radiation (0.9 Gy) with no rescue bone marrow. Results of control and transfected mice (control n = 5, *MNX1*: n = 8, *ETV6* and *MNX1::ETV6* n = 3) were analyzed using the log-rank test (ns: non-significant) (e). Data represents mean ± SD of at least three experiments and is considered significant using two-sided student t-test (**) at *p* ≤ 0.01 and (*) at *p* ≤ 0.05, or non-significant (ns) at *p* > 0.05. FL: Fetal liver, BM: bone marrow, ABM: Adult Bone Marrow, Ctrl: empty vector control, Fusion: *MNX1::ETV6* fusion, HGB: Hemoglobin, LSK: Lin^−^Sca-1^+^c-Kit^+^, NSG: NOD.Cg-*Kit*^*W-41J*^ *Tyr* ^+^ *Prkdc*^*scid*^ *Il2rg*^*tm1Wjl*^/ThomJ (NBSGW), H&E: hematoxylin and eosin. GFP: Green fluorescence protein, YFP: Yellow fluorescence protein.

### MNX1 alters differentiation in favor of myeloid lineage while increasing proliferation and colony replating capacity of the cells

In order to characterize this leukemia model, *in vitro* FL-HSPC (r-FL) transduced with *MNX1*::*ETV6, MNX1, ETV6* or empty retroviral vector control were assessed for their immunophenotype. Both *MNX1* and *MNX1*::*ETV6* altered differentiation in favor of myeloid lineage, with *MNX1* showing the most prominent effects. *MNX1* increased Mac-1 and Ly6C+ cells, accompanied by depletion of the Lin-/Sca1+/c-kit+ (LSK) population, while *MNX1*::*ETV6* only increased Ly6C+ cells (Figure 2a-b; Supplement figure 4b-d). Ectopic expression of MNX1 reduced the progenitor MEP population without significantly affecting CMP or GMP (Supplement figure 5a). Additionally, *MNX1* increased GEMM colonies with a concomitant reduction in BFU colonies and increased both CFU replating and proliferation capacity (Figure 2c-d; Supplement figure 6f-h). In ABM-HSPC *in vitro* cells (r-ABM), *MNX1* had similar effects but to a lower extent, and *MNX1*::*ETV6* fusion had no effects (Supplement figure 6 c-e).

**Figure 2.**
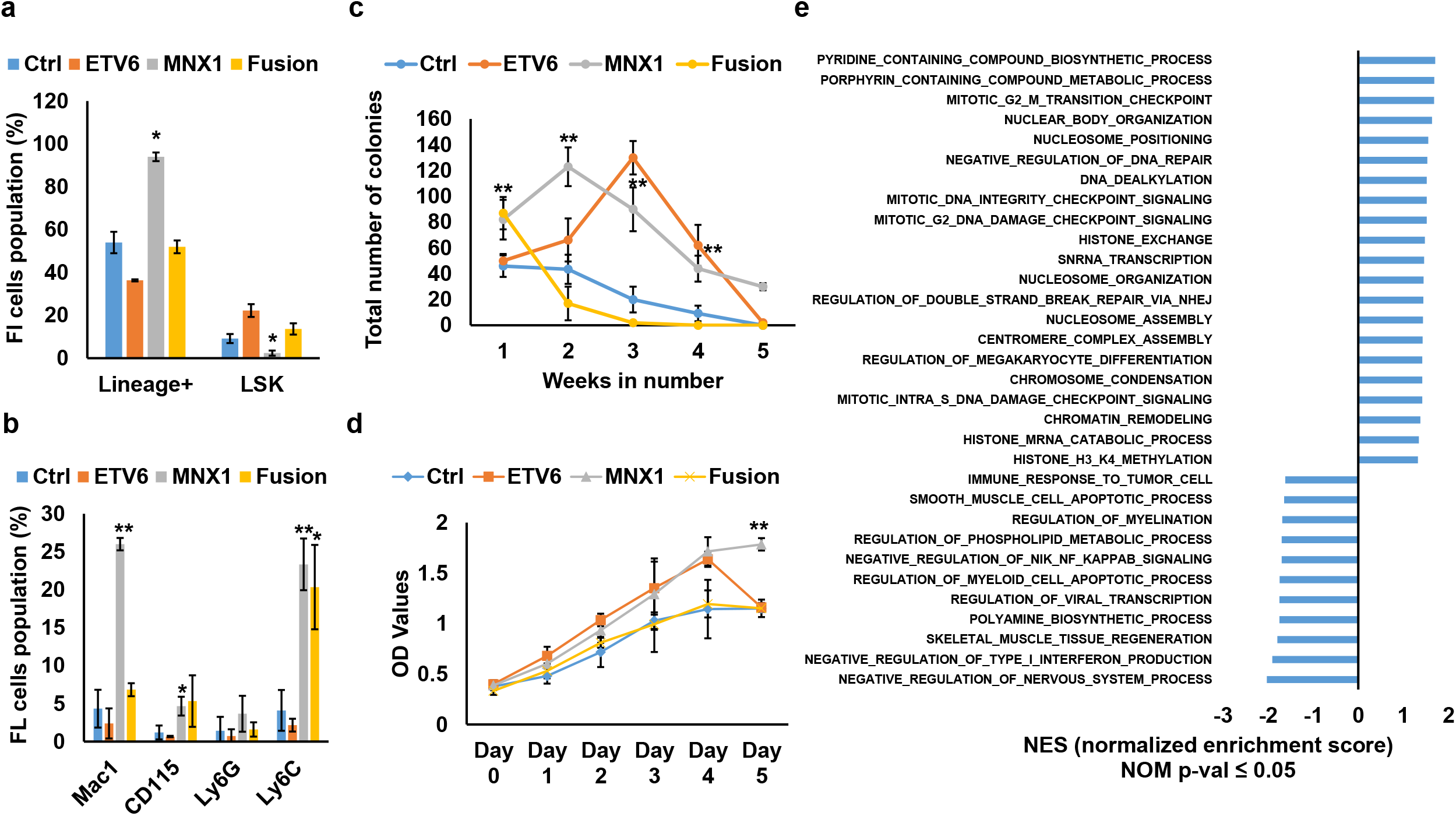
MNX1 alters differentiation and proliferation of FL-HSPC. Quantification of the flowcytometry analysis of *in vitro* FL-HSPC (r-FL) transduced with ectopic expression of *MNX1, ETV6, MNX1::ETV6* fusion or empty vector control with the indicated antibodies represented as percentage of population (a-b). Number of CFU colonies after transduction of FL-HSPC with replating for five consecutive weeks (c). MTT proliferation assay of transduced *invitro* FL-HSPC (r-FL) cells (d). Gene set enrichment analysis (GSEA) using the Gene ontology biological pathways (GO) gene set showing normalized enrichment score for pathways with NOM p-value ≤ 0.05 from leukemia BM cells with *MNX1* ectopic expression in comparison with FL-HSPC with empty vector control (e). Data represents mean ± SD of at least three experiments and is considered significant using two-sided student t-test (**) at *p* ≤ 0.01 and (*) at *p* ≤ 0.05. HSPC: hematopoietic stem and progenitor cells, FL: fetal liver, Ctrl: empty vector control, Fusion: *MNX1::ETV6* fusion, LSK: Lin^−^Sca-1^+^c-Kit^+^, lin+: lineage positive, r-FL: retroviral transduced *in vitro* FL cells, CFU: colony forming unit.

### MNX1 induces DNA damage

To investigate the molecular pathway through which *MNX1* is mediating its leukemogenic effect, differential gene expression between *MNX1* FL-HSCP leukemic NSG BM cells and FL-HSPC transduced with empty vector (Ctrl) was assessed with RNA-sequencing (Figure 2e Supplement figure 7). Gene ontology biological (GO) pathway and gene set enrichment (GSEA) analyses revealed that the highest enriched pathways in *MNX1* cells involved DNA damage, cell cycle, chromatin organization, methylation of histones, metabolic processes, megakaryocyte, and myeloid cell differentiation pathways (Figure 2e). Similar results were seen when using BM taken from mice transplanted with FL-HSCP with empty vector as control (Supplement figure 8). The effect on DNA damage was confirmed through examination of γH2AX foci induction. *MNX1* and to a lower extent *MNX1::ETV6* induced a higher number of γH2AX foci, indicative of higher DNA damage, in *in vitro* FL-HSPC as well as in ABM-HSPC (r-Fl & r-ABM) (Figure 3a-b; Supplement figure 9a-b). In both *MNX1* and *MNX1*::*ETV6* transduced FL-HSPC (r-FL), the DNA damage was accompanied with a transient G1 cell cycle accumulation and fewer cells in S-phase (Figure 3c; Supplement figure 9c). No such effect was observed in ABM-HSPC (r-ABM), where the G1 cells were replaced by a Sub-G1 peak suggestive of apoptotic cells after 3-4 weeks of transduction (Supplement figure 9d). Using Annexin-V and DAPI staining a 3.5-4-fold increase in apoptosis was induced by *MNX1* in ABM-HSPC (r-ABM), but not seen in FL-HSPC (r-FL) (Figure 3d-e). Following up the consequences of such pronounced DNA damage, we showed that BM from NSG leukemia mice exhibited an increase in the amount of DNA or a hyperploidy in comparison with the BM from Ctrl mice as indicated by increased DNA index (Supplement figure 10a-b). An interesting candidate for mediating DNA damage identified in several differentially regulated pathways was the centrosomal protein 164 (*Cep164*) (Supplement table 2). Increased expression of *Cep164* was confirmed with qPCR in both *MNX1* transduced FL-HSPC and leukemia BM (Supplement figure 10c). Furthermore, MNX1 increased binding to the promoter of *Cep164* was detected by ChIP-qPCR (Supplement figure 10c), along with altered binding of histone modifications that can change CEP164 expression namely H3K4me3 and H3K27me3 in the same promoter region (Supplement figure 10d).

**Figure 3.**
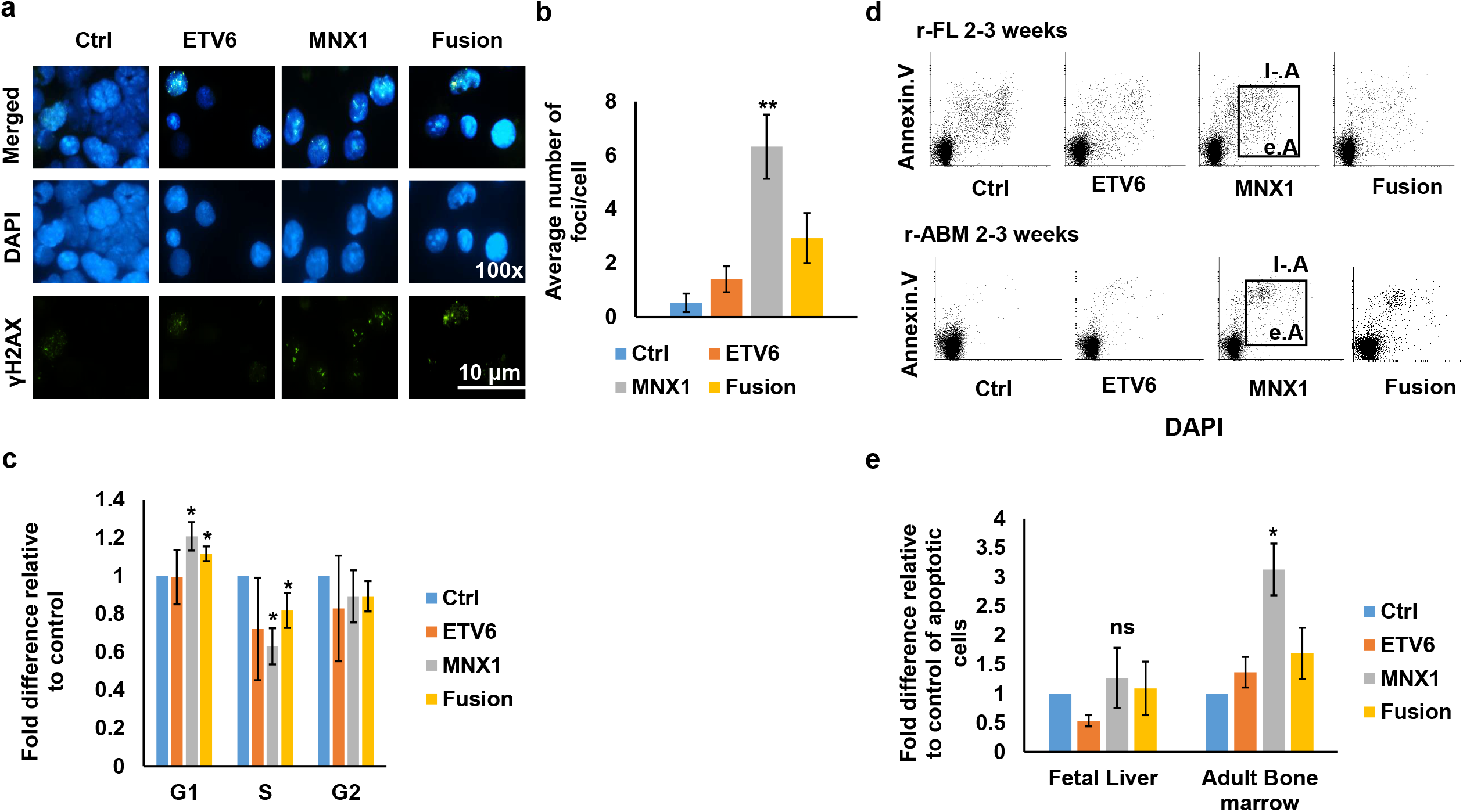
MNX1 induces DNA damage. Immunofluorescence of *in vitro* FL-HSPC (r-FL) cells stained with γH2AX –FITC antibody (green) and counterstained with DAPI (blue) (a). Quantification of the number of γH2AX foci/cell. At least 50 cells were counted (b). Quantification of cell cycle distribution from flowcytometry analysis represented as fold difference relative to control (c). Representative dot plots from the flow cytometry analysis of FL-HSPC (r-FL) and ABM-HSPC (r-ABM) after double staining with Annexin/V and DAPI for apoptotic analysis (d). Quantification of the flowcytometry analysis represented as fold difference relative to control (e). Data represents mean ± SD of at least three experiments and is considered significant using two-sided student t-test (**) at *p* ≤ 0.01 and (*) at *p* ≤ 0.05. FL: fetal liver, ABM: adult bone marrow, Ctrl: empty vector control, Fusion: *MNX1::ETV6* fusion, r-FL: retroviral *invitro* FL cells, r-BM: retroviral *in vitro* ABM cells e.A: early apoptosis, l-.A: late apoptosis.

### MNX1 alters histone modifications globally affecting chromatin accessibility

Since MNX1 induced histone modifications H3K4me3 and H3K27me3 at the *Cep164* promoter, we assessed global histone modifications induced by MNX1. Western blotting showed that MNX1 altered H3K4me3, H3K27me3, and both mono- and di-methylation of the H3K4 *in vitro* and in BM from NSG leukemic mice (Figure 4a; Supplement figure 10e). Co-immunoprecipitation experiments (Co-IP) confirmed the association of MNX1 to histone H3 (Supplement figure 11a). Antibody-guided chromatin tagmentation sequencing (ACT-Seq) was then used for mapping genome-wide distribution of the histone modifications (Figure 4b). H3K27me3 histone modification showed fewer accessible regions in MNX1 (BM from NSG leukemic mice) compared to Ctrl (FL-HSPC transduced with empty vector) (FDR ≤ 0.05, log2 fold change ≥ |1|), involving mainly distal intergenic regions followed by promoter and other intronic regions (Figure 4b). The consequences of these histone modifications on chromatin accessibility were investigated with ATAC-Seq. MNX1 leukemic BM cells exhibited increased number of differentially accessible chromatin regions in comparison with Ctrl FL-HSPC (FDR ≤ 0.05, log2 fold change ≥ |1|) (Figure 4c), mainly involving promoters, followed by distal intergenic and intronic regions, very similar to the pattern seen for H3K27me3 (Figure 4b). Pathway analysis of genes annotated to the differentially accessible regions from ATAC-Seq (FDR ≤ 0.05, log2 fold change ≥ |1|) revealed similar pathway enrichment to the RNA-Seq, with high enrichment in metabolic pathways, myeloid cell differentiation, erythrocyte differentiation, cytoskeleton organization, cell cycle process and apoptotic cell process (Supplement figure 11b). Further analysis through integrating ATAC-Seq data with in-silico predicted transcription factor binding sites by DiffTF package, revealed 130 differentially activated sites (FDR < 0.05) between MNX1 leukemic BM cells and Ctrl FL-HSPC (Table 1, Supplement figure 11c). These were enriched in pathways of myeloid/erythrocyte differentiation, cellular metabolism, G1/S-phase transition of cell cycle, histone methylation, DNA methylation and DNA damage response (Table1, Supplement figure 11c). A significant correlation between the accessible chromatin regions identified by ATAC-Seq and differentially expressed genes by RNA-Seq was shown at r_*S*_=0.47 and (p ≤ 0.01) (Figure 4d). In conclusion, MNX1 induces global histone modifications that are affecting chromatin accessibility and inducing differential gene regulation.

**Table 1.**
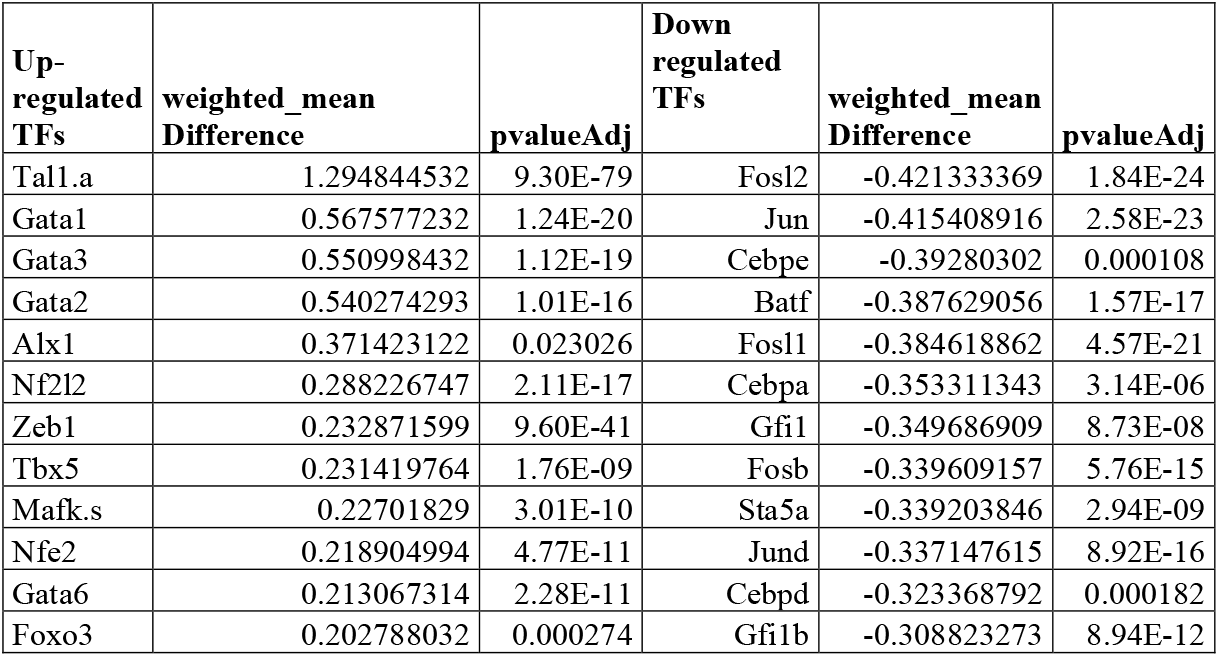
In-silico predicted transcription factor binding sites by DiffTF package from ATAC-Seq showing top differentially activated (FDR < 0.05) transcription factors (TFs) binding between MNX1 BM from leukemia mice and FL-HSPC Ctrl cells

| Up-regulated TFs | weighted_mean Difference | pvalueAdj | Down regulated TFs | weighted_mean Difference | pvalueAdj |
| --- | --- | --- | --- | --- | --- |
| Tal1.a | 1.294844532 | 9.30E-79 | Fosl2 | -0.421333369 | 1.84E-24 |
| Gata1 | 0.567577232 | 1.24E-20 | Jun | -0.415408916 | 2.58E-23 |
| Gata3 | 0.550998432 | 1.12E-19 | Cebpe | -0.39280302 | 0.000108 |
| Gata2 | 0.540274293 | 1.01E-16 | Batf | -0.387629056 | 1.57E-17 |
| Alx1 | 0.371423122 | 0.023026 | Fosl1 | -0.384618862 | 4.57E-21 |
| Nf2l2 | 0.288226747 | 2.11E-17 | Cebpa | -0.353311343 | 3.14E-06 |
| Zeb1 | 0.232871599 | 9.60E-41 | Gfi1 | -0.349686909 | 8.73E-08 |
| Tbx5 | 0.231419764 | 1.76E-09 | Fosb | -0.339609157 | 5.76E-15 |
| Mafk.s | 0.22701829 | 3.01E-10 | Sta5a | -0.339203846 | 2.94E-09 |
| Nfe2 | 0.218904994 | 4.77E-11 | Jund | -0.337147615 | 8.92E-16 |
| Gata6 | 0.213067314 | 2.28E-11 | Cebpd | -0.323368792 | 0.000182 |
| Foxo3 | 0.202788032 | 0.000274 | Gfi1b | -0.308823273 | 8.94E-12 |

**Figure 4.**
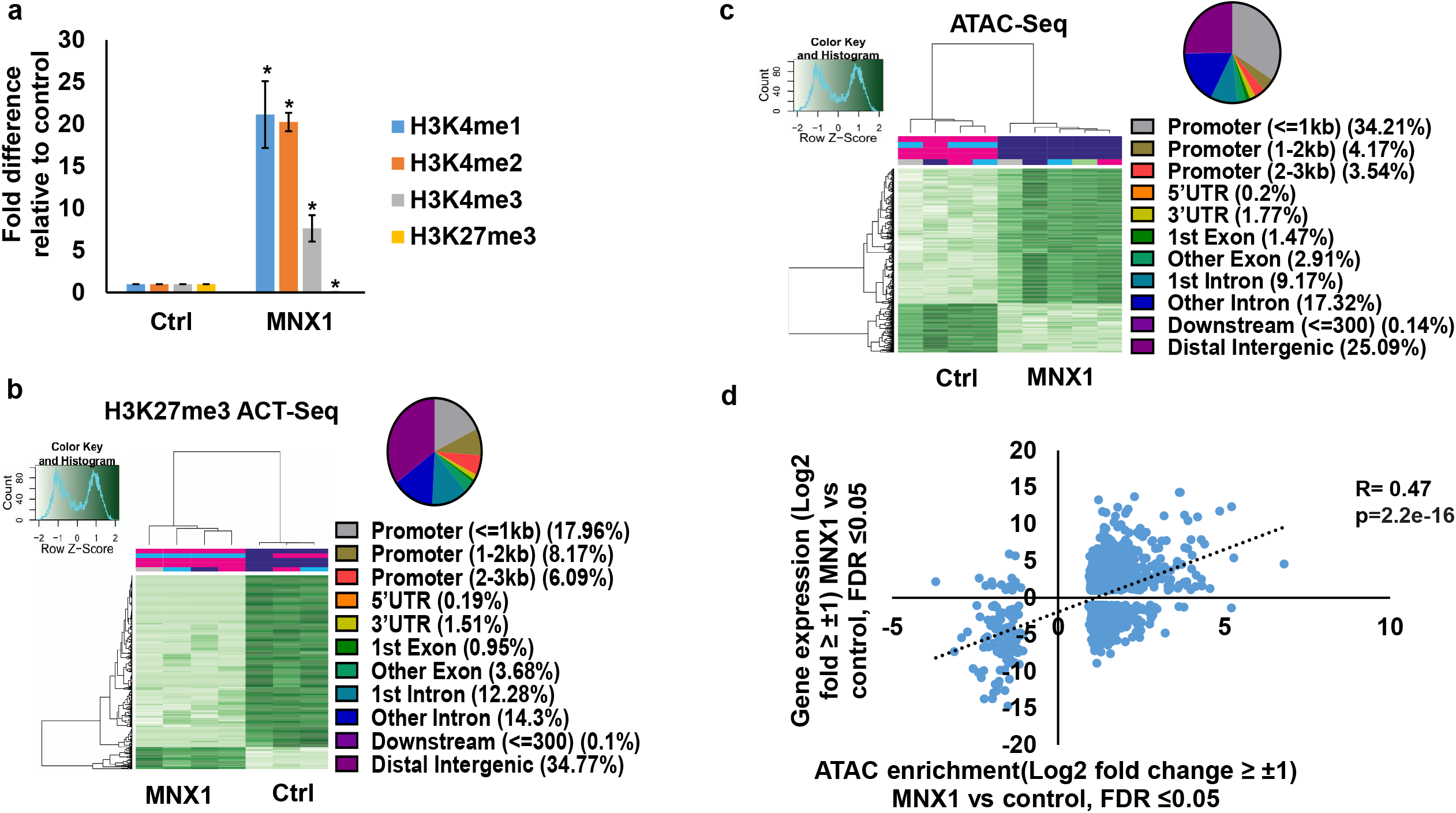
MNX1 alters histone methylation. Image quantification of western blot analysis of H3K4me1, H3K4me2, H3K4me3 and H3K27me3 as fold difference relative to loading control (a). Heatmap of differentially bound H3K27me3 from BM of leukemic mice (MNX1) versus control FL-HSPC transduced with empty vector (Ctrl) as determined by ACT-Seq and annotation of different enriched regions for H3K27me3 ACT-Seq (b). Results were considered at the log fold-change cutoff (logFC) of ≥ |1| and false discovery rate (FDR) of ≤0.05. Heatmap of differentially accessible regions from BM of leukemic mice (MNX1) versus control FL-HSPC transduced with empty vector (Ctrl) as determined by ATAC-Seq and annotation of different enriched regions for ATAC-Seq (c). Results were considered at the log fold-change cutoff (logFC) of ≥ |1| and false discovery rate (FDR) of ≤0.05. Scatter plot showing the correlation between differentially expressed genes from RNA-Seq expression analysis and genes with differentially accessible regions from ATAC-Seq analysis (d). Results were considered at the log fold-change cutoff (logFC) of ≥ |1| and false discovery rate (FDR) of ≤0.05.

### MNX1 alters the methylation of histone H3

To understand the mechanism for the MNX1-induced global histone modifications, we performed mass spectrometry analysis, to study proteins in association with MNX1. MNX1 associated proteins were pulled down after Co-IP from *in vitro* cells with anti-HA antibody. Pathway analysis using STRING for protein-protein interactions for the identified proteins revealed a high enrichment for methylation pathway proteins MAT2A, MAT2B and AHCY, (Figure 5a, Table 2), in addition to several S-adenosylmethionine (SAM) dependent methyl transferases and their downstream targets (Supplement table 3). Co-IP confirmed the association of MNX1 to MAT2A and AHCY both *in vitro* and in NSG BM leukemic cells (Figure 5b; Supplement figure 12a), with no effect on protein expression as shown in parallel with western blot (Supplement figure 12b). Furthermore, MNX1 overexpression increased the concentration of S-adenosylhomocysterin (SAH) and reduced free methionine in both FL cells and leukemia NSG BM cells (Supplement figure 12c-e). In support of a MNX1 role in methylation pathway and in altering histone methylation, MNX1 pulled down with Co-IP and incubated with recombinant Histone H3 resulted in methylation of Histone H3 (Figure 5c-d, Supplement figure 12f-g).

**Table 2.**
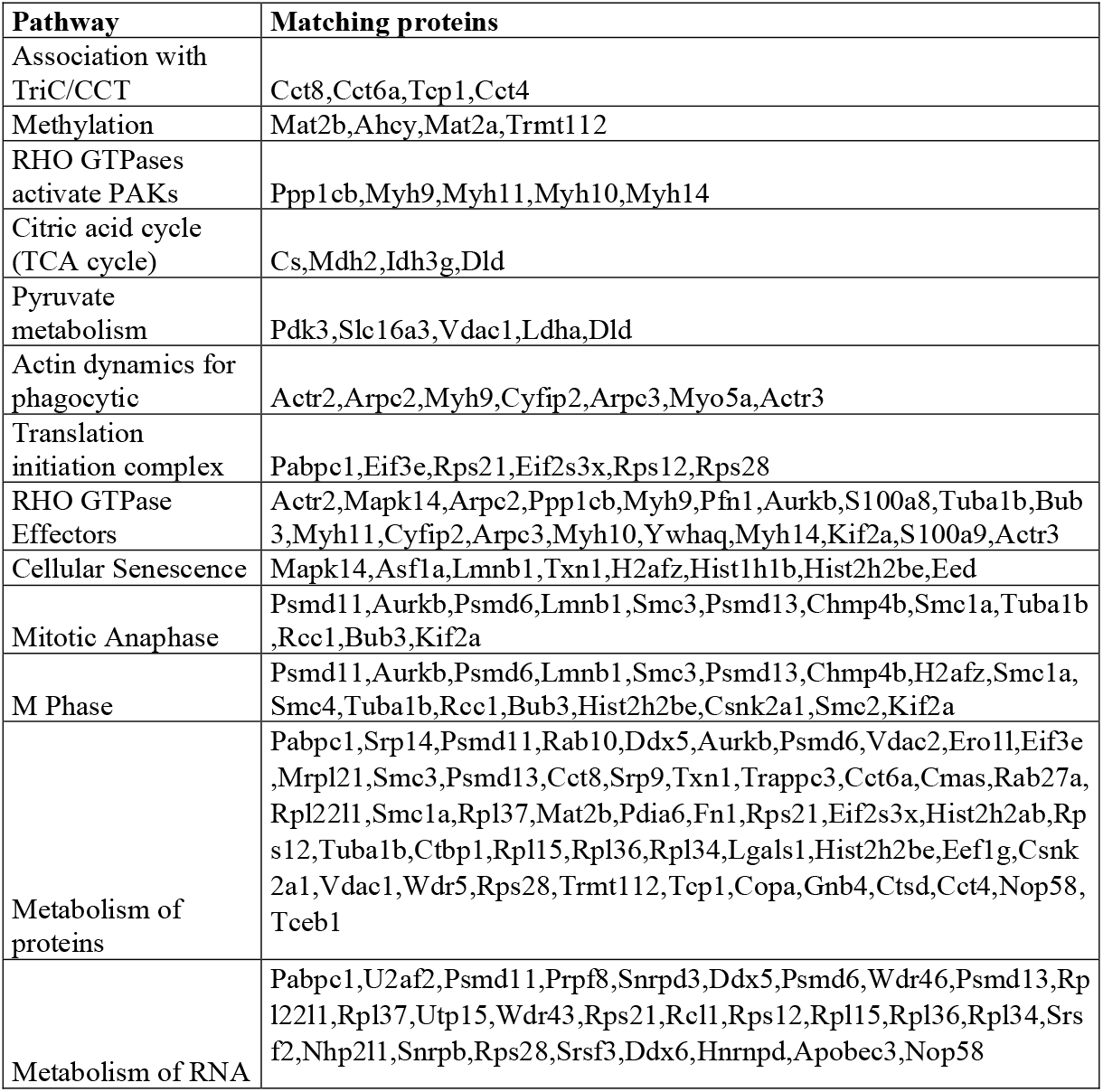
Identified proteins in top enriched pathways after Co-IP of proteins in complex with MNX1 and mass spectrometry analysis. As determined by STRING using reactome data set

**Figure 5.**
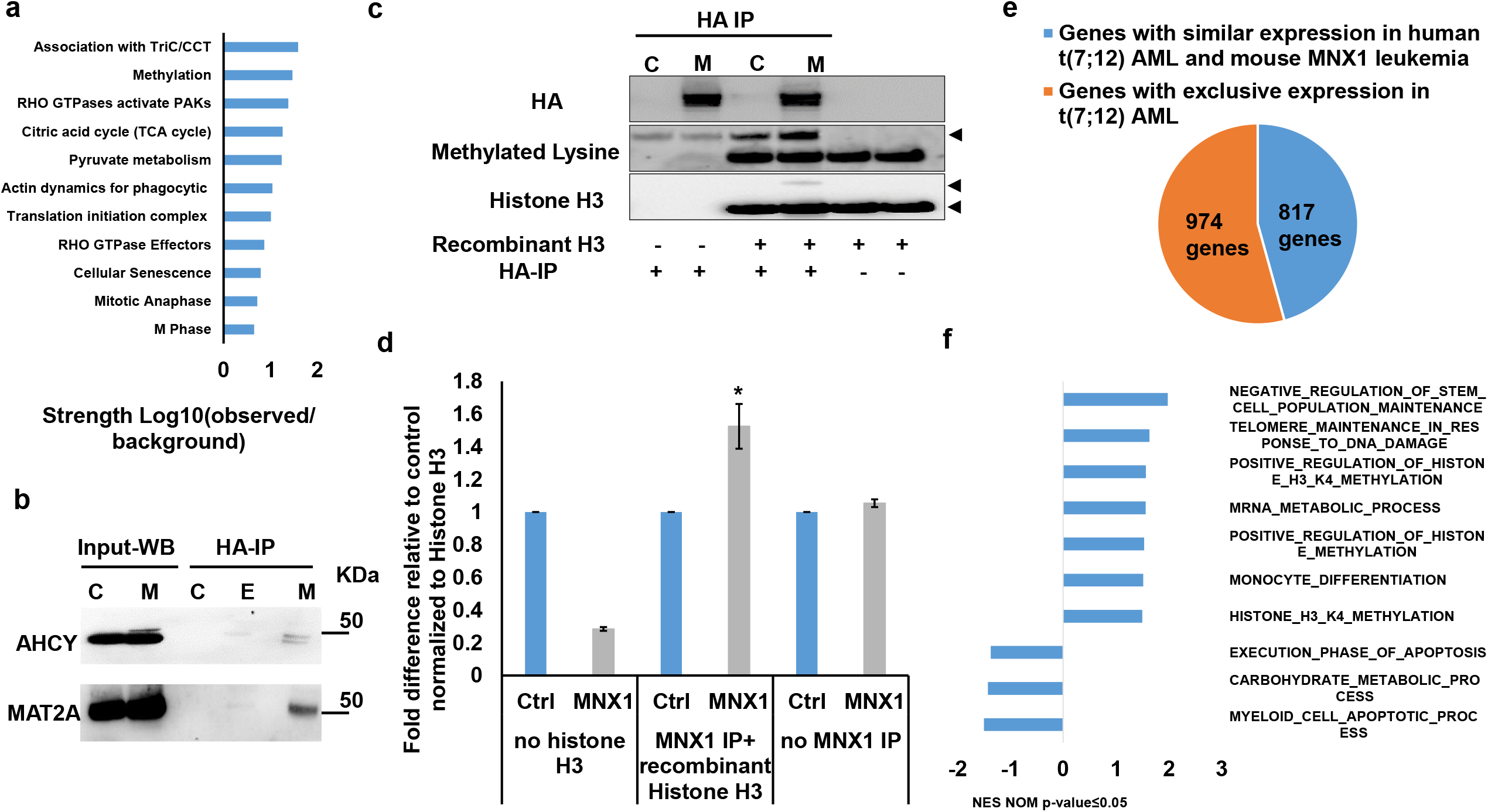
MNX1 associates with proteins from the methionine cycle and show similar gene expression and pathway enrichment with human t(7;12) AML. Pathway enrichment analysis for the identified proteins after Co-IP and mass spectrometry using STRING with reactome data set, showing strength of the pathway as Log10 observed proteins/background in the respective pathway (a). The binding of AHCY and MAT2A was detected by immunoprecipitation of MNX1 using HA-antibody followed by western blot using AHCY and MAT2A antibodies. ETV6 with HA tag was used as a negative control. Total protein input used as loading control (b). Recombinant Histone H3.1and H3.3 were subjected to an in vitro methyltransferase reaction using MNX1 complex pulled down with HA antibody from FL-HSPC transduced with MNX1(M), in comparison with FL-HSPC with empty vector control (C), in the presence of SAM and DTT. The reactions were terminated by boiling in SDS sample buffer. Separation of samples in 12% SDS-PAGE was followed by immunoblotting (IB) with mono, di and tri methyl-lysines antibody. Reblotting was made for detection of HA and total Histone H3. As indicated, negative controls were obtained by omitting Histone H3, or the pulled down immune complex (c). Quantification of the in vitro methyltransferase reaction (d). Data represents mean ± SD of at least three experiments and is considered significant using two-sided student t-test (**) at *p* ≤ 0.01 and (*) at *p* ≤ 0.05. Graph showing differentially expressed genes with same or opposite regulation when comparing genes expression data from t(7;12) AML patients from the TARGET database and RNA-seq data from BM from mice with MNX1 induced leukemia. Differentially expressed genes were selected based on a LogFC ±1 and FDR ≤ 0.05 (e). Gene set enrichment analysis (GSEA) using the Gene ontology biological pathways (GO), showing normalized enrichment score for common pathways between t(7;12) AML patients and the MNX1 induced leukemia in mice with NOM p-value ≤ 0.05 (b). r-FL: fetal liver, Ctrl: empty vector control, WB: Western blot, IP: immunoprecipitation. SAM: S-Adenosyl methionine, DTT: Dithiothreitol.

### MNX1 induced leukemia in mice and human pediatric t(7;12) AML show similar gene expression and pathway enrichment

To validate the similarity between the AML developed in our mouse model with human t(7;12) AML, we used our RNA-sequencing data on leukemic cells from mice and retrieved RNA-sequencing data from pediatric t(7;12) AML patient samples from the children’s oncology group (COG)–National Cancer Institute (NCI) TARGET AML initiative data set. Differential gene expression in mouse leukemia was determined by comparing *MNX1* FL-HSCP leukemic NSG BM cells and FL-HSPC transduced with empty vector and differential gene expression in human AML was determined by comparing pediatric t(7;12) AML patient samples with normal human BM. Comparing the differential gene expression between the mouse AML with MNX1 expression and the t(7;12) patient data revealed close to 50% overlapping differential gene expression (Figure 5e; Supplement table 4). These included increased expression of the genes *MNX1, cKIT, CEP164, AHCY, MAT2A* and *MAT2B* (Supplement figure 13). Pathway enrichment analysis of the t(7;12) patient using gene set enrichment (GSEA) analyses revealed several common enriched pathways including; DNA damage, H3K4methylation, monocyte differentiation, apoptotic processes and metabolic processes (Figure 5f and supplement figure 14). Interestingly, only the H3K4me3 methylation, and no other histone methylation pathway was enriched in the GSEA analysis from t(7;12) AML patients. Thus, our data highlight the biological significance for MNX1 expression in pediatric t(7;12) AML and suggesting a mechanism where MNX1 is mediating its oncogenic effects through aberrant methylation resulting in histone modifications resulting in leukemia transformation.

### The effects of MNX1 on methylation are crucial for leukemogenesis

To confirm the importance of this effect for MNX1-induced leukemia, we used the natural nucleoside analogue of SAM, Sinefungin, which acts as a competitor and accordingly a pan-methyltranferase inhibitor^17^. Adding 5μM of Sinefungin to MNX1-transduced FL-HSPC partially or completely prevented the effects of MNX1 on histone modifications (Figure 6a, Supplement figure 15a), DNA damage induction (Figure 6b; Supplement figure 15b), cell cycle distribution (Supplement figure 15c), myeloid differentiation and LSK depletion (Figure 6c-d). Furthermore, when MNX1 FL-HSPC was pretreated *in vitro* with Sinefungin and then transplanted into NBSGW mice there were no signs of leukemia development (Figure 6e; Supplement figure 15d-e), despite maintained high MNX1 expression (Supplement figure 15f) and continuous presence of viable transplanted cells in blood (Supplement figure 16a) and. To investigate the effect of Sinefungin treatment on MNX1 induced differential gene expression, RNA sequencing was done before and after treatment. Differentially expressed genes (FDR ≤ 0.05, log2 fold change ≥ |1.5|) from MNX1 cells after treatment (MNX+S) clustered with MNX1 (Figure 6f), and most of the differentially expressed genes by MNX1 overexpression remained at similar level after treatment (Figure 6f), with limited number of genes altered in MNX1 differential gene expression after treatment (Supplement figure 12b)

**Figure 6.**
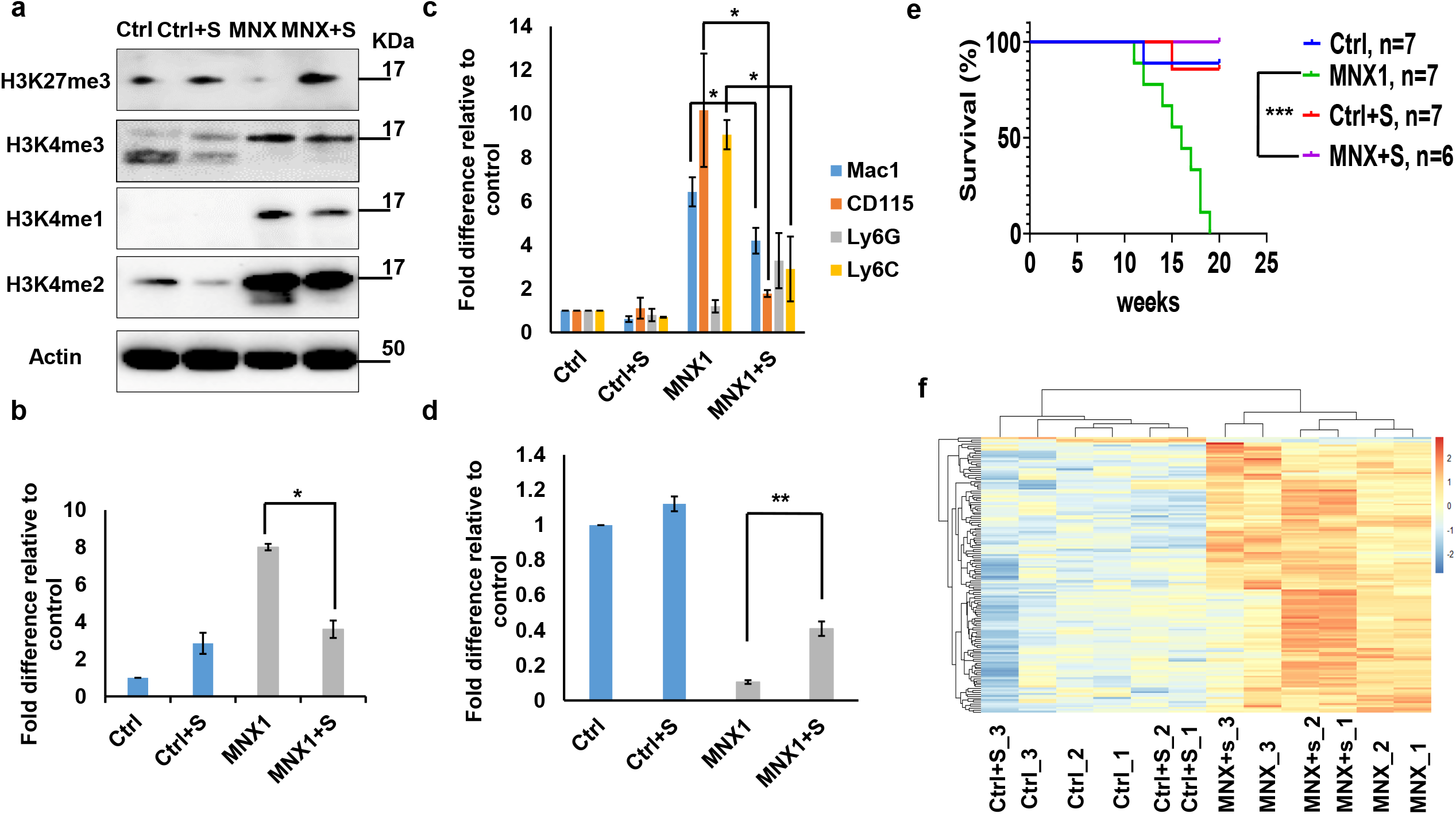
Sinefungin rescued MNX1-induced phenotype. Protein expression levels of H3K4me3, H3k27me3, H3K4me1 and H3K4me2 in *invitro* FL-HSPC (r-FL) cells with either MNX1 overexpression or empty vector (Ctrl) after treatment with vehicle or 5μM Sinefungin (Ctrl+S, MNX1+S) as determined by Western blotting. Actin served as loading control (a). Quantification of the Western blot analysis in Supplement figure 9a. Quantification of the number of γH2AX foci/cell. At least 50 cells were counted. Data represented as fold difference relative to control (b). Quantification of flow cytometry analysis of the cells with the indicated antibodies. Data represented as fold difference relative to control (c and d). Data represents mean ± SD of at least three experiments and is considered significant using two-sided student t-test between MNX1 and MNX1+S (**) at *p* ≤ 0.01 and (*) at *p* ≤ 0.05. Kaplan-Meier survival curves of (NBSGW) mice transplanted with FL cells after retrovirus transduction with either ectopic expression of MNX1 or empty vector (Ctrl) after treatment with vehicle or 5μM Sinefungin (MNX1+S, Ctrl+S). Results of MNX1 (n = 7) and Sinefungin treated MNX1 cells (MNX1+S) (n = 6) were analyzed using the log-rank test (***: significant at *P* ≤ 0.01) (e). LogFC heat map of downregulated (blue) and upregulated (red) differentially expressed genes of FL cells with *MNX1* ectopic expression (MNX) in comparison with FL cells with empty vector (Ctrl), with 5μM Sinefungin (Ctrl+S, MNX+S) or without treatment (Ctrl, MNX1), showing clustering and similarity between the samples. Results were considered at the log fold-change cutoff (LogFC) ≥ |1.5| and FDR ≤ 0.05 (f).

## DISCUSSION

The t(7;12) has only been reported in children diagnosed with AML before the age of 24 months. The function of this translocation in inducing infant leukemia and the reason for its absence in adult leukemia is unknown. In the current study using a murine model, we showed that ectopic expression of MNX1 rather than the *MNX1*::*ETV6* fusion was able to initiate and drive leukemogenesis. Our data suggest a mechanism through which MNX1 is mediating the leukemogenic effect through aberrant methylation that results in histone modifications and DNA damage.

The malignant transformation mediated by MNX1 overexpression in our mouse model matched the criteria for AML ^18^, compatible with an AML without maturation. This mouse MNX1 driven leukemia had a high degree of differentially expressed gene that overlap with the genes gene expression signature and pathway enrichment that is seen in human AML with t(7;12). It has also been shown that MNX1 clearly has oncogenic properties in both human and murine hematopoietic cells by inducing a myeloid biased perturbed hematopoietic differentiation and premature senescence ^19^. This fits well with the properties seen in our mouse leukemia model with both block in differentiation and induced cell cycle arrest. Similar gene expression and pathway enrichment has also been shown between human cells with an engineered t(7;12) translocation, which results in high expression of MNX1, and human t(7;12) AML which suggests a common gene expression program induced by MNX1 in hematopoietic cells ^20, 21^. Our data showed that ectopic expression of MNX1 was able to induce AML using HSPC from fetal origin but not from adult bone marrow. One possible reason for this was the dramatic induction of apoptosis seen in the hematopoietic progenitor cells from adult bone marrow, prohibiting leukemic transformation. The higher susceptibility for apoptosis and DNA damage induced by MNX1 is concordant with the presence of naturally occurring DNA damage in the adult stem cells ^22^. Possibly, the balance between fetal and adult stem cell programs affects the transforming ability of the cells upon overexpression of MNX1. Lin28b has been shown as a key regulator of the self-renewing capacity characteristic of fetal but not adult hematopoietic stem cells ^23^. *LIN28B* is, together with *MNX1*, a signature gene expressed in all pediatric t(7;12) AML and not seen in other AML subtypes ^24^. Lin28b was seen expressed in our MNX1 induced mouse leukemia (Supplement table 2), which suggests that transformation by MNX1 might be dependent on Lin28b for its transforming effects or perhaps more likely indicates that the fetal hematopoietic program is needed for MNX1 oncogenesis. However, the expression of *Lin28b* in our leukemia and *LIN28B* in human t(7;12) AML is contradictory to the finding that Lin28b suppresses MLL-ENL fusion driven leukemogenesis, also typically seen in infants leukemia ^25^. But what is intriguing is that the MLL-ENL fusion’s potential to initiate leukemia development peaks during neonatal development and drops dramatically in adult mice ^25^. The importance of the development stage of cells that can be transformed into leukemia including the leukemia phenotype as well as intrinsic properties of progenitor populations have been shown in several studies^26-28^. Other factors that differ between fetal and adult hematopoetic stem cells and that might affect their propensity for transformation are metabolic demand and cell cycle profile ^29-31^. Even though MNX1 overexpression induced leukemia in the cells of fetal origin, the fusion *MNX1*::*ETV6* by itself did not induce leukemia. This finding is in line with the previously reported inability to induce transformation with *MNX1*::*ETV6* (*HLXB9/TEL*) *in vitro* and the paucity of transgenic mouse models of AML with *MNX1*::*ETV6* ^32^. In our study, the development of leukemia induced by MNX1 expression was primarily seen in immunocompromised NSG mice. Thus, the adaptive B- and T-cell immune response might be enough to eradicate cells with overexpression of MNX1 and prevent leukemia development. This may be yet another clue to the development of AML with t(7;12) in very young children, typically before six months of age when the immune system is still under development ^4-6^.

Our studies of MNX1 expression *in vitro* revealed an induction of DNA damage both in FL-HSPC and ABM-HSPC, which could contribute to the observed skewed differentiation towards myeloid lineage evident by the depletion of LSK and MEP population and increased Mac1+ and Ly6C+ population. The influence of DNA damage in stem cells on differentiation was first demonstrated in melanocyte stem cells, where ionizing radiation triggered differentiation into mature melanocytes ^33^. In hematopoietic stem cells, DNA damage can induce differentiation towards lymphoid or myeloid lineage and may be the reason for the skewing towards myeoloid differentiation in the aging hematopoietic system ^34-37^.

We found MNX1 to associate with members of the methionine cycle, including MAT2A, AHCY and MAT2B, in addition to several downstream SAM-dependent methyl transferases. Methionine is an essential amino acid that is converted to the universal methyl donor SAM, which upon the donation of its methyl group is converted to SAH. This reaction is catalyzed by methionine adenosyl transferases (MAT). SAM is used as a cofactor in most methylation reactions and provides the activated methyl group for methylation of proteins, including histones, DNA, RNA and lipids. These methylation events are highly dependent on methionine metabolism, with alterations in methionine showing profound effects on DNA and histone methylation ^38, 39^. The role of another homeodomain protein (MSX1) in recruiting methyltransferase to regulate gene expression and chromatin structure through histone modifications has been in the differentiation of myoblasts ^40, 41^. During neural development, MNX1 binding to loci on chromatin are enriched for H3K4me1 and H3K4me3 ^42^. Therefore, the binding and association of MNX1 with methyltranferases and members of methionine cycle, and the subsequent change in chromatin structure and histone modifications might be the physiological role for MNX1 during differentiation ^43, 44^. We conclude that the abnormal expression of MNX1 and subsequent effect on the methionine cycle and chromatin structure in hematopoietic cells acts as the driver of leukemia transformation, supported by the inhibition of the phenotype by the SAM analog Sinefungin.

In conclusion, our results provide the biological and clinical significance for MNX1 as an epigenetic regulator in pediatric t(7;12) AML. Given that many epigenetic modifications are chemically reversible, the inhibition of MNX1 ectopic expression or its downstream effects in t(7;12) could provide a foundation for alternative treatment options to improve outcome.

## Acknowledgment

We thank Mohamad Ali and Akram Mendez for help with bioinformatic analysis. Tova Johansson and Hanna Brissman for help with animals. This work was supported by grants from the Swedish Cancer Society (CAN2017/461), the Swedish Childhood Cancer Foundation (PR2014-0125, PR2019-0013 and TJ2019-0053, TJ2022-0017), Wilhelm och Martina Lundgrens Fond, Assar Gabrielsson Fond and Västra Götalandsregionen (ALFGBG-431881). The German Funding Agency (DFG) with funding for Collaborative Research Center 1074, Project B11N to CP. The computations were enabled by resources in project [SNIC 2021/22-754] provided by the Swedish National Infrastructure for Computing (SNIC) at UPPMAX, partially funded by the Swedish Research Council through grant agreement no. 2018-05973.

