## Supplemental material for "Aberrant MNX1 Expression Associated with t(7;12)(q36;p13) Pediatric Acute Myeloid Leukemia Induces the Disease Through Altering Histone Methylation"

***ABSTRACT***

Certain subtypes of acute myeloid leukemia (AML) in children have inferior outcome. One of these has a translocation t(7;12)(q36;p13) leading to a *MNX1*::*ETV6* fusion along with high expression of MNX1. Here we identified the transforming event in this AML and possible ways of treatment. Only MNX1 was able to induce AML in mice, with similar gene expression and pathway enrichment to t(7;12) AML patient data. Importantly, this leukemia was only induced in immune incompetent mice using fetal but not adult hematopoietic stem and progenitor cells. The restriction in transforming capacity to cells from fetal liver is in alignment with t(7;12)(q36;p13) AML being mostly seen in infants. Expression of MNX1 led to increased histone 3 lysine 4 mono-, di- and trimethylation, reduction in H3K27me3, accompanied with changes in genome-wide chromatin accessibility and genome expression, likely mediated through MNX1 interaction with the methionine cycle and methyltransferases. MNX1 expression increased DNA damage, depletion of the Lin-/Sca1+/c-Kit+ population and skewing toward the myeloid lineage. These effects, together with leukemia development, was prevented by pretreatment with the S-adenosylmethionine analog Sinefungin. In conclusion, we have shown the importance of MNX1 in development of AML with t(7;12), supporting a rationale for targeting MNX1 and downstream pathways.

#### Supplementary materials and methods

##### *Western blotting*

Cells were lysed in modified RIPA buffer (50 mm Tris, pH 7.4, 150 mm NaCl, 1% Nonidet P-40, 1 mm EDTA, 0.25% sodium deoxycholate) containing protease (Roche Applied Science, Mannheim, Germany) and phosphatase (Sigma) inhibitors and processed as described previously [1]. Cell lysate subjected to sonication using bioruptor for 5 minutes (30 second ON/ 30 seconds OFF). Primary antibodies used in this study included the following: rabbit-anti HA # 11846, rabbit-anti actin # 4970, rabbit-anti H3K4me1 #5326, rabbit-anti H3K4me2 #9725 from Cell Signaling Technology, rabbit-anti Mat2A#NB110-94158, rabbit-anti Ahcy #NBP2-48817 from Novous biological, rabbit-anti H3K4me3 (catalog no. C15410003-50), rabbit anti-H3K27me3 (catalog no. C15410069), from Diagenode and rabbit anti-Histone 3 (catalog no. Ab18521) from Abcam.

##### *Immunoprecipitation and mass spectrometry*

Anti HA-magnetic beads from Thermo Fisher Scientific (catalog no. 88836) were used for immunoprecipitation. After blocking with 0.1% BSA in PBS, 0.7-1 mg of cell lysates were incubated with HA antibody for 1h at RT. For IgG antibody, the antibody was cross-linked to magnetic protein G Dynabeads using Dynabeads® antibody coupling kit 14311D (Invitrogen). The immune complexes were washed three times with lysis buffer and eluted by boiling in SDS sample buffer (Invitrogen). Precipitates were separated with SDS-PAGE and co-precipitated protein bands were visualized using Coomassie Blue. Prominent gel bands were excised for LC-MS using a hybrid LTQ-Orbitrap Velos mass spectrometer (Thermo Fisher Scientific) as described in [1].

***Invitro histone methyltransferase***

Invitro histone methyltransferase was done as described in [2]. Briefly, 0.25-0.5 mg of cell lysates were incubated with HA antibody for 1h at RT. The immune complexes were washed three times with lysis buffer and once with BC100 buffer (20 mM Tris–HCl pH 8.0, 100 mM KCl, 0.2 mM EDTA, 20% glycerol, 1 mM DTT). The immune complexes was incubated with 4 μg of human recombinant histone H3.1 and H3.3 (New England biolabs # M2507S, # M2503S) at 30 °C for 60 min in 25 μl of methylation buffer buffer (50 mM Tris–HCl pH 8.5, 5 mM MgCl2) in the presence of 10 μM SAM (New England biolabs # B9003S) and 4mM DTT. Omission of the recombinant histone H3, or the immune complexes served as negative controls. The reactions were terminated by boiling in SDS sample buffer (Invitrogen), followed by separations in 12% SDS-PAGE and immunoblottings using H3k4m31 (Cell signaling), Histone H3 (Abcam) and anti-methlylated lysines (Abcam #ab23366). Quantification was done using ImageJ [3].

***Immunophenotyping and flowcytometry***

Cells were collected 1-2 weeks after transduction. Then they were washed twice with PBS, blocked with 0.1% FBS, mouse BD FC Block #553142 (BD Biosciences) in PBS for 10 mins RT. Then incubated with the antibodies for 20 minutes at RT, washed and resuspended in PBS. The following antibodies were purchased from BD Biosciences. Sca1-V450#560653, Cd3e-V450#560801, Ter119-V450#560504, CD45R-V450#560473, Ly6G/Ly6C-V450#560454, Mac1-V450#560455, Ly6G-PeCy7#560601, B220-V450#560472, CD127-V450#561205, cKit-APC#553356, Sca1-PeCy7#558162, CD45-APCCy7#561037, Ter119-APC#561033, CD8a-APCCy7#561967, CD4-APCCy7#561830. The following from (ThermoFisher Scientific). CD115-APC#17-1152-80, CD64-APCCy7#47-0641-82.

For cell cycle analysis, the proportion of cells in $G_1$ and $G_2$ phases were measured by staining with DAPI (BD Biosciences, #564901) as described previously (41). Briefly, cells were washed with PBS, and stained DAPI solution 1:100 containing RNase A (250 μg/ml) and 0.25% Triton X100 for 15 min at room temperature. The samples were processed using FACSAria (BD Biosciences), and the data were analyzed with the Cell Quest software (BD Biosciences) or with flowing software v 2.5.1 (Turku bioscience center, Finland).

Apoptosis analysis was studied using Annexin V/DAPI method (Annexin V-FLUOS staining kit, Roche, Mannheim, Germany) according to the manufacturer's protocol using AnexinV-APC antibody#550474 (BD Biosciences) and DAPI. Flow cytometric analysis was immediately performed using FACSAria.

***Histological preparation***

Moribund mice or mice at the end of the experiment were euthanized. Liver, spleen, and sternum were harvested by fixing in 4% formaldehyde at 4°C, dehydrated in 70–100% ethanol, cleared in xylene and embedded in paraffin. Sections (4–5 μm). Paraffin-embedded tissue sections were prepared at Histocentre (Gothenburg, Sweden) and stained with HandE. Blood samples was collected directly from the heart after euthanizing the mice , prepared as thin blood films, Giemsa-stained and examined under the microscope (Nikon Inc., Japan).

***Colony forming unit assay***

Methocult M3434 (STEMCELL Technologies) was used for the CFU assays where 5000 cells were seeded/petri dish and incubated for 7-14 days according to the manufacturer’s protocol.

***Proliferation assay***

Cell proliferation was determined using MTT (3-[4,5-dimethylthiazol-2-yl]-2,5-diphenyltetrazolium bromide) assay from abcam # ab211091. Briefly, FL cells transducted with our constructs were plated, in triplicate, in a 96-well culture plate at 37° C in CO2 incubator for 7 days. Each day an equal volume of MTT was added to the cell culture medium. The plate was incubated at 37° C for 4 h followed by addition of MTT solvent. The plate was then swirled gently until all crystals are dissolved. Absorbance values were blanked against cell culture medium without cells. The absorbance was read at 590 nm using a microplate reader (*Infinite F50®Tecan* Group Ltd., Swiss). The absorbance was plotted against the number of days

***Sinefungin treatment***

Sinefungin was purchased from Calbiochem #567051 as Insolution. Final doses of 5µM was used to treat the cells. Daily change of medium with fresh Sinefungin was added after viral transfection for one week.

***Gene expression analysis***

Total RNA was isolated using the miRNeasy Plus micro kit (Qiagen) and cDNA synthesized with SuperScript III First-Strand Synthesis SuperMix for qRT-PCR (Invitrogen/Thermo Fisher Scientific). TaqMan Gene Expression Assays and TaqMan Universal MasterMix II (Applied Biosystems/ Thermo Fisher Scientific) were used for probe based qPCR assays, listed in supplement table 1. A SYBR Green assay was used for the MNX1, ETV6 and MNX1(ex1)::ETV6(ex3) fusion detection, with primers synthetized at IDT (Integrated DNA Technologies) and Power SYBR Green PCR Master Mix (Applied Biosystems/Thermo Fisher Scientific), listed in supplement table 1. The reactions were performed on the ABI 7900HT Sequence Detection System (Applied Biosystems/ Thermo Fisher Scientific) with *hprt* as the reference gene. Gene expression was calculated using the deltaCt method and are presented as $2^{-deltaCt}$.

***Next-generation sequencing***

Total RNA was extracted from BM cells from mice with MNX1 induced leukemia and from transduced FL cells after one week of transfection as control using RepliePrep RNA miniprep systems (Promega). 5-20µg of total RNA were utilized for RNA-Seq using DNBseq technology at Beijing Genomics Institute (BGI, Hong Kong), with RIN value of ≥7.5 and 100bp paired-end reads. An average of 40 million reads were generated for each sample. The gene expression level was quantified by BGI Europe. Only samples with 28S/18S ≥1, Q20_rate ≥ 97 and GC% ≥ 51% were used. For RNA-seq analysis, alignment was carried out to reference mouse genome GRCm38 using STAR (version 2.9.0a) with the following parameters: outSAMtype BAM SortedByCoordinate. Quantification was carried out using HTSeq-count (Python package HTSeq, python v 3.8.1) software using the following parameters -f bam -r pos -s no -m union with the reference annotation file. The mouse reference genome and annotation files were obtained from the Ensembl database. Differential gene expression was performed using the R package "Deseq2" [4]. Genes with log2 (foldchange) 1 and false recovery rate (FDR) adjusted p-value < 0.05 were considered differentially expressed genes. Principal component (PCA) analysis, heatmap analysis (pheatmap) and enhanced volcano plot (enhancedVolcano) were performed in order to examine the level of differences in the gene expression profiles between control and MNX1 samples. Gene Ontology Go-biological process database was used for pathway analysis using the default parameters. A pathway was considered to be significantly enriched for a FDR < 0.05, p-value < 0.05 of the respective pathway. Gene Set Enrichment Analysis (GSEA) was used with the Gene ontology GO-biological process gene set. A pathway in GSEA analysis was regarded significant at nominal p-value < 0.05.

***Assay for transposase-accessible chromatin using sequencing (ATACseq)***

ATACseq libraries were prepared according to a modified Omni-ATAC protocol [5] as follows. Cells stored at -80 °C in culture medium containing 10% DMSO were thawed on ice, counted with a Countess counter (Invitrogen), pelleted at 1000 × g for 10 min at 4 °C, and a suitable aliquot for both ATACseq and antibody-guided chromatin tagmentation (ACTseq) was resuspended in cold phosphate-buffered saline (PBS) at a titer of $10^6$ viable cells/ml. An aliquot of 50000 viable cells was pelleted at 600 × g for 5 min at 4 °C, and cells were resuspended in 50 µl cold RSB containing each 0.1% of NP40 and Tween-20 and 0.01% digitonin. Proper lysis was monitored microscopically using Trypan blue, then 200 µl RSB containing 0.1% Tween-20 were mixed in. Lysed cells were spun with 1000 x g for 10 min at 4 °C and resuspended in 48 µl transposition buffer (10 mM Tris-acetate, pH 7.6, 5 mM $MgCl_2$, 10% dimethylformamide, 30% PBS, 0.01% digitonin and 0.1% Tween-20). The suspension was completed by addition of 2 µl Tn*5* transposome generated as described by Picelli et al. [6]. (The Tn*5* transposase was kindly provided by the EMBL-Heidelberg protein expression and purification core facility). Tagmentation was performed at 37 °C for 30 minutes in a thermomixer with 1000 rpm. DNA was purified after addition of 20 µl 5 M guanidinium-thiocyanate with 140 µl HighPrep beads (Biozym) and eluted with 20 µl $H_2O$. Sequencing libraries were generated by addition of 25 µl NEBNext High Fidelity 2× Master Mix (NEB), each 2.5 µl of 10 µM primer Tn5mCP1n and Tn5mCBar [7] and 0.5 µl of 100x SYBRGreen I using a LightCycler 480 real-time machine with 72 °C for 5 min, 98 °C for 30 sec and cycling at 98 °C, 10 sec; 63 °C, 30 sec; 72 °C, 30 sec. Amplification was stopped when an increase of about 10 fluorescence units was observed, usually between cycle 12 and 15. . Libraries were bead-purified with 70 µl HighPrep beads and eluted with 12 µl EB (Qiagen). Concentrations and fragment lengths were determined with the Qubit dsDNA High sensitivity kit (Invitrogen) and the TapeStation D1000 High sensitivity assay (Agilent), respectively. Up to ten libraries with compatible barcodes were pooled equimolarly, and pools of 5-10 nM were sequenced in the DKFZ Genomics and Proteomics Core Facility (GPCF) using an Illumina NextSeq 550 system with 75 bp paired-end, mid output mode. Sequencing reads were bioinformatically analyzed as detailed by Hey et al. [8]. In short, peak calling of processed reads was done using MACS2 callpeak v. 2.1.0.20140616 with a q-value cutoff of 0.05 and default parameters. The analysis is a publicly accessible fully containerized workflow using Common Workflow Language v. 1.0 (https://doi.org/10.6084/m9.figshare.3115156.v2); Highly Portable Processing of Data Generated by ChIP-seq, ChIPmentation, Cut&Run, and ACT-seq. Verion 1.1.2, Zenodo: Temple, FL, USA). Differentially accessible regions (DARs) were identified with the DiffBind R package (R package version 2.12.0) [9]. Common peaks were considered those found in at least two samples. Differential analysis was performed using edgeR [10]. Regions with an FDR < 0.05 were considered differentially accessible. Further annotation of DARs was performed with R package ChIPseeker (R package version 1.20.0) [11]. Differential transcription factor activity was predicted using diffTF [12] and in silico predicted transcription factor binding sites based on the HOCOMOCO 10 database were used as a reference [13].

***Antibody-guided chromatin tagmentation followed by sequencing (ACTseq)***

ACTseq [14] was essentially performed as detailed by Liu et al. [15]. As described in the ATACseq methods chapter, frozen cells were thawed, washed with PBS and, per antibody, aliquots of 50000 viable cells were lysed. Antibodies used for the pA-Tn*5* transposome-antibody complexes were directed against H3K27me3 (Cell signaling, 9733), IgG (Millipore PP64B) and yeast histone H2B (M3930, Boster Bio. Tech., Pleasanton, CA, USA), the latter complexes being used for spike-in to enable sequence read normalization between biological replicates [15]. Sequencing libraries were generated, sequenced and bioinformatically analyzed as described in the ATACseq section. For data processing, Trim Galore v. 0.4.4[16] (https://www.bioinformatics.babraham.ac.uk/projects/trim_galore/ ; accessed on 13 December

2020) was applied together with Cutadapt v. 1.14 [17] using “–paired”, “–nextera”, “– length_1 35”, and “–length_2 35” parameters. Trimmed reads were mapped against the Genome Reference Consortium Human Build version 37 by means of Bowtie2 v. 2.2.6 [18] using the “–very-sensitive” flag and a maximum insertion length of 2500 bp. Mappings belonging to the same lane-multiplexed library were combined using SAMtools merge v. 1.5 [19]. Discordant alignments and mappings with a Phred score below 20 were removed using SAMtools [20], as fragments obtained from tagmentation cannot be smaller than 38 bp. Thus, all alignments corresponding to fragment sizes below that threshold were removed. Read ends were shifted to represent the center of the transposition event as previously described by [21]. Additionally, trimmed reads were aligned against the Saccharomyces cerevisiae R64 reference genome followed by post-alignment filtering as described above. To calculate a library-specific scaling factor, we derived the multiplicative inverse of the number of filtered reads mapped to the yeast  genome. Coverage tracks were generated by means of the bamCoverage functionality of Deeptools v. 3.1.1 [22] using the non-default parameters “–ignoreForNormalization chrM chrY chrX” and “–effectiveGenomeSize 2652783500” as well as the “–scaleRatio” option to specify the spike-in-based scaling factor. The analysis procedure was implemented as fully containerized workflow using the Common Workflow Language v. 1.0. Differentially bound regions were identified as described for the ATACseq data.

***Chromatin immunoprecipitation (ChIP):***

ChIP assays were performed using iDeal ChIP-qPCR kit (C01010180, Diagenode). Cells were subject to fixation and crosslinking by formaldehyde (net concentration 1%) for 10-15 minutes at RT. Formaldehyde fixation is quenched by adding Glycine 125uM and incubating the fixed cellular suspension at RT for 5-10 minutes. The pellet is then washed twice by cold PBS and subjected then to lysis according to the specifications of the manufacturer. Next, cell lysates is subjected to sonication using bioruptor for 20 minutes (30 second ON/ 30 seconds OFF), so that the sheared smear is supposed to range from 200-600 bp long. The sheared chromatin is then subject for immunoprecipitation using high quality ChIP-grade antibodies against: HA (cell signaling), H3K27me3 (Diagenode), H3K4me3 (Diagenode), and polyclonal rabbit anti-IgG (c15410206, Diagenode) as negative control. Immunoprecipitation relies on that each antibody is crosslinked to protein A/G-coated magnetic beads, and the reaction takes up to 16 hours at 4$^{o}$C, and then the beads crosslinked with the antibody-DNA is then subject for washing by buffers containing gradient concentrations of detergent. The washed beads-antibody-DNA is then subjected for de-crosslinking in the presence of proteinase K, so that all DNA-binding proteins, histones and beads are released, and the DNA sequences are then eluted and used for downstream ChIP-real time PCR. The final eluted DNA was analyzed by real-time quantitative PCR with the Power SYBR Green PCR master mix (Applied Biosystems). The PCR conditions were: 50 °C for 2 min, 95 °C for 10 min, and 40 cycles of 95 °C for 15 s and 60 °C for 1 min. The PCR analysis was performed with the 7900HT fast real-time PCR system instrument and software (Applied Biosystems). All primers used to amplify the genomic target sites were designed by using PRIMER3 PLUS online tools. Primer sequences are listed in Supplement table 2.

The calculation of enrichment was normalized to the respective inputs and done according to the following formulas:

• The % of IP to input formula: % recovery = 2^[(Ct input-log2(5%)) – Ct sample] * 100%

• The enrichment was calculated as: %recovery IP / % recovery IgG

Negative control was achieved by immunoprecipitation against HA antibody, followed by detection of random region with low CpG islands and promoter binding density corresponding to chr1:168430579+168430704

***Methionine concentration:***

Amino acid quantification was done at the Swedish Metabolomics Centre, Umeå, Sweden by LC-MSMS. This was done as following:

Standards and Calibration Curve; Amino acid standards (alanine, arginine, aspartic acid, cysteine, glutamic acid, glycine, histidine, isoleucine, leucine, lysine, methionine, phenylalanine, proline, serine, threonine, tyrosine, valine, glutamine, asparagine, GABA, citrulline, ornithine, taurine, tryphtophan, 5-HTP and kynurenine) were purchased from Sigma (St. Louis, MO, USA). Isotopically labeled amino acid standards (alanine ($^{13}C_3$, $^{15}N$), arginine ($^{13}C_6$, $^{15}N_4$), aspartic acid ($^{13}C_4$, $^{15}N$), cystine ($^{13}C_6$, $^{15}N_2$), glutamic acid ($^{13}C_5$, $^{15}N$), glycine ($^{13}C_2$, $^{15}N$), histidine ($^{13}C_6$, $^{15}N_3$), isoleucine ($^{13}C_6$, $^{15}N$), leucine ($^{13}C_6$, $^{15}N$), lysine ($^{13}C_6$, $^{15}N_2$), methionine ($^{13}C_5$, $^{15}N$), phenylalanine ($^{13}C_9$, $^{15}N$), proline ($^{13}C_5$, $^{15}N$), serine ($^{13}C_3$, $^{15}N$), threonine ($^{13}C_4$, $^{15}N$), tyrosine ($^{13}C_9$, $^{15}N$), valine ($^{13}C_5$, $^{15}N$), Citrulline (d4), GABA ($^{13}C_4$), glutamine ($^{13}C_5$), asparagine($^{13}C_4$), ornithine (d6), tryptophan (d8), kynurenine (d4) ) were obtained from Cambridge Isotope Laboratories (Andover, MA). Stock solutions of each compound were prepared at a concentration of 500 ng/µL and stored at -80ºC. A 10-point calibration curve (0.01-100 pmol/µL) was prepared by serial dilutions and spiked with internal standards at a final concentration of 5 pmol/µL. Mass spectrometry grade formic acid was purchased from Sigma-Aldrich (St Louis, MO, USA) and HPLC grade acetonitrile from Fisher Scientific (Fair Lawn, NJ, USA). T

***Amino acid extraction***

One million cells were extracted with 400µL 90:10 (v/v) Methanol:water solution containing internal standards at 2.5 nmol/µL. One tungsten bead were added and the samples were shaken at 30 Hz for 2 min in a mixer mill. After removing the bead, the samples were centrifuged at 4 °C, 14000 RPM, for 10 min. The supernatant collected and stored at -80 °C until analysis.

*Amino acid derivatization with AccQ-Tag*

Extracted samples were derivatized by AccQ-Tag™ (Waters, Milford, MA, USA) according to the manufacturer's instructions. Briefly, 10 µL of extract were added to 70 µL of AccQ•Tag Ultra Borate buffer and finally 20 µL of the freshly prepared AccQ•Tag derivatization solution was added, and the sample was immediately vortex for 30s. Samples were kept at room temperature for 30 minutes followed by 10 minutes at 55ºC. For each batch quality control samples and procedure blanks were included. Calibration curves were prepared in a similar way by adding 10 µL of each point of the curve to 70 µL of AccQ•Tag Ultra Borate buffer and 20 µL of the freshly prepared AccQ•Tag derivatization solution.

*Amino acids Quantification by LC-ESI-MSMS*

Derivatized solutions were analyzed using a 1290 Infinitely system from Agilent Technologies (Waldbronn, Germany), consisting G4220A binary pump, G1316C thermostated column compartment and G4226A autosampler with G1330B autosampler thermostat coupled to an Agilent 6460 triple quadrupole mass spectrometer equipped with a jet stream electrospray source operating in positive ion mode. Separation was achieved injecting 1 µL of each sample onto a BEH $C_{18}$ 2.1x100 mm, 1.7 µm column (Waters, Milford, MA, USA) held at 50ºC in a column oven. The gradient eluents used were $H_2O$ 0.1% formic acid (A) and acetonitrile 0.1% formic acid (B) with a flow rate of 500 µL/min. The initial conditions consisted of 0% B, and the following gradient was used with linear increments: 0.54-3.50 minutes (0.1-9.1% B), 3.50-7.0 (9.1-17.0% B), 7.0-8.0 (17.0-19.70% B), 8.0-8.5 (19.7% B), 8.5-9.0 (19.7-21.2% B), 9.0-10.0 (21.2-59.6% B), 10.0-11.0 (59.6-95.0% B), 11.0-11.5 (95.0% B), 11.5-15.0 (0% B). From 13.0 minutes to 14.8 minutes the flow rate was set at 800 µL/min for a faster equilibration of the column. The MS parameters were optimized for each compound as described in Supporting Information. MRM transitions for the derivatized amino acids were optimized using

MassHunter MS Optimizer software (Agilent Technologies Inc., Santa Clara, CA, USA). The fragmentor voltage was set at 380 V, the cell accelerator voltage at 7 V and the collision energies from 14-45V, nitrogen was used as collision gas. Jet-stream gas temperature was 290°C with a gas flow of 11 L/min, sheath gas temperature 325°C, sheath gas flow of 12 L/min. The nebulizer pressure was set to 20 psi and the capillary voltage was set at 4 kV. The QqQ was run in Dynamic MRM Mode with 2 min retention time windows and 500 msec cycle scans.

The data was quantified using MassHunter™ Quantitation software B08.00 (Agilent Technologies Inc., Santa Clara, CA, USA) and the amount of each amino acid was calculated based on the calibration curves.

***Retention times (rt), MRM-transition stages monitored (precursor ion and product ions) and collision energies of analyzed compounds.***

| Compounds | MRM transition | | rt (min) | Collision Energy (V) |
|---|---|---|---|---|
| | Precursor Ion | Product Ion | | |
| alanine | 260.1 | 171 | 4.1 | 14 |
| arginine | 345.1 | 171 | 2.8 | 35 |
| aspartic acid | 304.1 | 171 | 3.4 | 18 |
| cystine | 581 | 171 | 5.2 | 25 |
| glutamic acid | 318.1 | 171 | 3.6 | 22 |
| glycine | 246.1 | 171 | 3.1 | 18 |
| histidine | 326.1 | 171 | 2.6 | 26 |
| isoleucine | 302.1 | 171 | 7.8 | 18 |
| leucine | 302.1 | 171 | 7.6 | 18 |
| lysine | 487.2 | 171 | 5.1 | 26 |
| methionine | 320.1 | 171 | 6.0 | 22 |
| phenylalanine | 336.1 | 171 | 8.1 | 18 |
| proline | 286.1 | 171 | 4.4 | 14 |
| serine | 276.1 | 171 | 3.1 | 14 |
| threonine | 290.1 | 171 | 3.7 | 18 |
| tyrosine | 352.1 | 171 | 5.7 | 18 |

| | | | | |
|---|---|---|---|---|
| valine | 288.1 | 171 | 6.1 | 18 |
| citrulline | 346.2 | 171 | 3.4 | 30 |
| GABA | 274.1 | 171 | 4.1 | 18 |
| glutamine | 317.1 | 171 | 3.0 | 22 |
| asparagine | 303.1 | 171 | 2.8 | 18 |
| ornithine | 473.2 | 171 | 4.7 | 34 |
| tryptophan | 375.2 | 171 | 8.4 | 26 |
| kynurenin | 379.2 | 171 | 7.6 | 37 |
| Labelled internal standards | | | | |
| alanine (13C3, 15N) | 264.07 | 171 | 4.1 | 14 |
| arginine (13C6, 15N4) | 355.1 | 171 | 2.8 | 35 |
| aspartic acid (13C4, 15N) | 309.07 | 171 | 3.3 | 18 |
| cystine (13C6, 15N2) | 589 | 171 | 5.2 | 25 |
| glutamic acid (13C5, 15N) | 324.09 | 171 | 3.5 | 22 |
| glycine (13C2, 15N) | 249.05 | 171 | 3.1 | 18 |
| histidine (13C6, 15N3) | 335 | 171 | 2.6 | 26 |
| isoleucine (13C6, 15N) | 309.12 | 171 | 7.9 | 18 |
| leucine (13C6, 15N) | 309.12 | 171 | 7.6 | 18 |
| lysine (13C6, 15N2) | 495.1 | 171 | 5.2 | 26 |
| methionine (13C5, 15N) | 326.17 | 171 | 6.0 | 22 |
| phenylalanine (13C9, 15N) | 346 | 171 | 8.2 | 18 |
| proline (13C5, 15N) | 292.09 | 171 | 4.4 | 14 |
| serine (13C3, 15N) | 280.6 | 171 | 3.0 | 14 |
| threonine (13C4, 15N) | 295.08 | 171 | 3.7 | 18 |
| tyrosine (13C9, 15N) | 362.12 | 171 | 5.7 | 18 |
| valine (13C5, 15N) | 294.1 | 171 | 6.1 | 18 |
| Citrulline (d4) | 350.1 | 171 | 3.4 | 30 |
| GABA (13C4) | 278.1 | 171 | 4.0 | 18 |
| glutamine (13C5) | 322.1 | 171 | 3.1 | 22 |
| asparagine(13C4) | 307.1 | 171 | 2.8 | 18 |
| ornithine (d6) | 479 | 171 | 4.6 | 34 |
| tryptophan (d8) | 383.2 | 171 | 8.3 | 26 |

| | | | | |
|---|---|---|---|---|
| kynurenine (d4) | 383 | 171 | 7.7 | 37 |

##### *SAH concentration*

Intracellular metabolites were isolated after lysating the cells in modified ripa buffer. Cell lysate was subjected to sonication using bioruptor for 5 minutes (30 second ON/ 30 seconds OFF). Resultant free cell supernatant was snap frozen and stored at -80 °C. Protein concentrations were determined using BCA (Pierce Biotechnology, Rockford, IL) and equal amounts of protein were used for SAH quantification. Quantification of intracellular SAH was done using S-Adenosylmethionine (SAM) and S-Adenosylhomocysteine (SAH) ELISA Combo Kit # MET-5151-C (Cell biolabs, San Diego, CA, USA) as instructed.

##### *t(7;12) patients RNA-Seq analysis*

Count read files from twelve t(7;12) patients versus and 62 normal human BM were obtained from the children's oncology group (COG)–National Cancer Institute (NCI) TARGET AML initiative data set (available at https://target-data.nci.nih.gov/Public/AML/mRNA-seq/). Differential gene expression was performed using the R package “DeSeq2. Genes with log2 (foldchange) 1 and false recovery rate (FDR) adjusted p-value < 0.05 were considered differentially expressed genes. Gene expression values were obtained from the normalized counts of DeSeq2. Gene Set Enrichment Analysis (GSEA) was used with the Gene ontology GO-biological process gene set for pathway enrichment analysis. A pathway in GSEA analysis was regarded significant at nominal p-value < 0.05. Converting mouse to human genes for comparison between the t(7;12) patients RNA-Seq and our MNX1 mouse model was established using the mouse genome informatics, available at (http://www.informatics.jax.org/downloads/reports/HOM_MouseHumanSequence.rpt)

### Supplemental Table Legends

**Supplement Table 1**

Table of the *MNX1*, *ETV6* and *MNX1::ETV6* fusion gene sequences, primer sequences for their detection by RT-PCR, TaqMan assays used and primers used for ChIP-qPCR.

**Supplement Table 2**

Differential gene expression of leukemia BM cells with *MNX1* ectopic expression (MNX) in comparison with FL cells with empty vector (Ctrl). Differential gene expression was performed using the R package "Deseq2". Genes with log2 (foldchange) 1 and false recovery rate (FDR) adjusted p-value < 0.05 were selected for downstream analysis.

**Supplementary Table 3.** Identified proteins after MNX1 pull down with Co-immunoprecipitation. IgG was used as negative control. Experiment was repeated twice and the common proteins that was absent in the negative control were selected for downstream analysis.

**Supplementary Table 4.** Expression pattern between human t(7;12) AML and mouse MNX1 leukaemia RNA-Seq data. Human t(7;12) AML RNA-Seq data was obtained from the children's oncology group (COG)–National Cancer Institute (NCI) TARGET AML initiative data set. Converting mouse to human genes for comparison between the t(7;12) patients RNA-Seq and our MNX1 mouse model was established using the mouse genome informatics.

### Supplementary Figure Legends

**Supplementary figure 1.** mRNA expression of *MNX1, ETV6* and *MNX1::ETV6* fusion determined by quantitative real time qPCR, normalized to *Hprt* and represented as relative expression from both *invitro* FL cells (a) and *invitro* adult bone marrow cells (b). Western blot analysis of HA in the transfected FL cells. Actin was used as loading control (c). Expected size for MNX1 (41-48 kDa) and MNX1::ETV6 fusion (72 kDa).

**Supplementary figure 2.** Kaplan-Meier survival curves of (C57BL/6) mice transplanted with either FL (a) cells or ABM (c) cells after lethal radiation (8.5 Gy) with rescue bone marrow. Cells were retrovirally transfected with either ectopic expression of *MNX1, ETV6, MNX1::ETV6* fusion or empty vector (Ctrl). Mice were euthanized at the end of the experiment and were analyzed for spleen weight, white blood cell number and hemoglobin concentration (HGB) (b&d). Kaplan-Meier survival curves of (C57BL/6) mice transplanted with FL cells after sub-lethal radiation (5.5 Gy) with no rescue bone marrow (e). Mice were euthanized at the end of the experiment and were analyzed for spleen weight, white blood cell number and hemoglobin concentration (f). Results of control ($n \geq 6$) and transfected mice ($n \geq 6$) were analyzed using the log-rank test (ns: non-significant).

**Supplementary figure 3.** Percentage of GFP/YFP chimerism in peripheral blood of (C57BL/6) mice transplanted with FL cells after lethal radiation (8.5 Gy) with rescue bone marrow, as determined by flowcytometry every two weeks (a). Percentage of MNX1 engraftment in bone marrow of (C57BL/6) in mice transplanted after lethal radiation (8.5 Gy) with rescue bone marrow versus mice transplanted after sublethal radiation (5.5 Gy) with no rescue BM, as determined by GFP/YFP in bone marrow of the euthanized mice with flowcytometry (b). Quantification of flow cytometry analysis of BM from (C57BL/6) mice transplanted with FL cells after sub-lethal radiation (5.5 Gy) with no rescue bone marrow and the indicated antibodies (c). Quantification of flow cytometry analysis of BM from NOD.Cg-*Kit*$^{W-41J}$ *Tyr* $^{+}$ *Prkdc*$^{scid}$ *Il2rg*$^{tm1Wjl}$/ThomJ (NBSGW) mice transplanted with FL cells with no radiation and the indicated antibodies (d). Kaplan-Meier survival curves of NOD.Cg-*Kit*$^{W-41J}$ *Tyr* $^{+}$ *Prkdc*$^{scid}$ *Il2rg*$^{tm1Wjl}$/ThomJ (NBSGW) mice transplanted with *MNX1*, *MNX1::ETV6* fusion or empty vector (Ctrl) after lethal radiation (1.6 Gy) with rescue BM (e, left panel), or after sublethal radiation (0.9 Gy) with no rescue BM (e, middle, panel). Results of control (n ≥ 6) and transfected mice (n ≥ 3) were analyzed using the log-rank test (***: significant at $P \leq 0.01$). Mice from sublethal radiation experiment were euthanized at the end and were analyzed for spleen weight, white blood cell number and hemoglobin concentration (e, right panel). Data represents mean ± SD of at least three experiments and is considered significant using two-sided student t-test (**) at $p \leq 0.01$ and (*) at $p \leq 0.05$.

GFP: Green fluorescence protein, YFP: Yellow fluorescence protein

**Supplementary figure 4.** Gating strategy for BM from leukemia mice in Figure 1. Gating was set using forward- and side-scatters (FSC and SSC), to distinguish between viable cells from cell debris. CD45 and Lineage were used to gate for hematopoietic cells which are lineage negative only represented by CD45+/lineage- population. Sca1 and cKit were used to determine the stem cell population (LSK) from the CD45+/lineage- population, represented by Sca1+, cKit+, lineage-. Alternatively, the percentage of cKit+, Ly6c+, Mac1+, CD115+, Ly6g+, CD4/CD8a+, B220+ or Ter119+ population was determined from the CD45+ population representing the hematopoietic cells (a). Gating strategy for invitro FL cells (r-FL) cells in Figure 2. Gating was set using forward- and side-scatters (FSC and SSC), to distinguish between viable cells from cell debris. CD45, GFP/YFP and Lineage were used to gate for hematopoietic cells which are lineage negative, positive for the transduced constructs and positive for the hematopietic cells marker CD45; represented by CD45+/lineage-/GFP+ population. Sca1 and cKit were used to determine the stem cell population (LSK) from the

CD45+/lineage-/GFP+ population, represented by Sca1+, cKit+, lineage-. Alternatively, the percentage of Ly6c+, Mac1+, CD115+, Ly6g+, CD4/CD8a+, B220+ or Ter119+ population was determined from the CD45+/GFP+ population representing the hematopoietic cells with the transduced constructs (b-d).

GFP: Green fluorescence protein, YFP: Yellow fluorescence protein

**Supplementary figure 5.** Kaplan-Meier survival curves of (NBSGW) mice transplanted with either primary FL cells retrovirally transfected with *MNX1* (red) or secondary transplanted with BM from leukemia mice (purple) and no radiation (a). Kaplan-Meier survival curves of (NBSGW) mice secondary transplanted with BM after sublethal radiation (0.9 Gy) (b, left panel). Mice were euthanized at the end of the experiment and were analyzed for spleen weight, white blood cell number and hemoglobin concentration (b, right panel). Quantification of flow cytometry analysis of BM from (NBSGW) mice transplanted with ABM cells after sub-lethal radiation (0.9 Gy) and no rescue bone marrow with the indicated antibodies (c). Cells were retrovirally transfected with either ectopic expression of *MNX1*, *ETV6*, *MNX1::ETV6* fusion or empty vector (Ctrl). Mice were euthanized at the end of the experiment and were analyzed for spleen weight, white blood cell number and hemoglobin concentration (HGB) (d). Data represents mean ± SD of at least three experiments and is considered non-significant using two-sided student t-test (ns) at $p > 0.05$.

**Supplementary figure 6.** Quantification of flow cytometry analysis of *invitro* FL cells (r-FL) (a-b) or ABM (r-ABM) cells (b-e) transfected with ectopic expression of *MNX1*, *ETV6*, *MNX1::ETV6* fusion or empty vector with the indicated antibodies. Quantification of the number of colonies after two weeks of CFU (Colony forming unit assay) (f). Data represents mean ± SD of at least three experiments and is considered significant using two-sided student t-test (**) at $p \leq 0.01$ and (*) at $p \leq 0.05$. CFU of transfected FL cells showing different types of colonies after 2 weeks (g). Bright-field microscopy image of Fl cells transfected with *MNX1* or empty vector (Ctrl), showing number of cells after 5 days (h).

GEMM:, Granulocyte-erythroid macrophage megakaryocytic colony-forming units, GM:, Granulocyte macrophase colony-forming cells BFU: Erythrocyte burst forming unit

**Supplementary figure 7.** mRNA expression of *MNX1* determined by quantitative real time qPCR, normalized to *Hprt* and represented as relative expression from BM of leukemia mice (a). Data represents mean ± SD of at least three experiments and is considered significant using two-sided student t-test (**) at $p \leq 0.01$. LogFC heat map of downregulated (blue) and upregulated (red) differentially expressed genes of leukemia BM cells with *MNX1* ectopic expression (MNX) in comparison with FL cells with empty vector (Ctrl). Differentially expressed genes were selected based on a LogFC1 and FDR ≤ 0.05 (b). PCA scatter plot of normalized gene expression using variance Stabilizing Transformation (c). Volcano plot of the differentially expressed genes (d).

**Supplementary figure 8.** LogFC heat map of downregulated (blue) and upregulated (red) differentially expressed genes of leukemia BM cells with *MNX1* ectopic expression (MNX) in comparison with BM from mice transplanted with FL with empty vector (Ctrl) (a). Differentially expressed genes were selected based on a LogFC1 and FDR ≤ 0.05. PCA scatter plot of normalized gene expression using variance Stabilizing Transformation (b). Gene set enrichment analysis (GSEA) using the Gene ontology biological pathways (GO), showing normalized enrichment score for pathways with NOM p-value ≤ 0.05 (c)

**Supplementary figure 9.** Immunofluorescence of *invitro* ABM cells (r-ABM) stained with γH2AX –FITC antibody (green) and counterstained with DAPI (blue) (a). Quantification of the number of γH2AX foci/cell. At least 50 cells were counted (b). Flowcytometry analysis of *invitro* FL (r-FL) and ABM (r-ABM) cells after staining with DAPI for cell cycle distribution (c-d). Quantification of the flowcytometry analysis for cell cycle distribution using ABM cells represented as fold difference relative to control (c, right panel). Data represents mean ± SD of at least three experiments and is considered significant using two-sided student t-test (*) at $p \leq 0.05$.

**Supplementary figure 10.** Flowcytometry analysis for cell cycle distribution showing the median DNA values at G1 phase and G2 phase of cell cycle for flowcytometry analysis of BM from leukemia NBSGW mice transfected with MNX1-FL cells in comparison with mice transfected with empty vector Ctrl (a-b). mRNA expression of *Cep164* determined by quantitative real time qPCR, normalized to *Hprt* and represented as fold difference relative to control from both in vitro FL-HSPC (MNX1-FL) and BM from leukemia mice (MNX1-BM) (c, left panel). Quantification of the binding of MNX1 to the promoter regions of *Cep164* in FL-HSPC with either MNX1 or empty ctrl determined by ChIP-qPCR (c, right panel). Quantification of the binding of H3K4me3 and H3K27me3 to the promoter regions of *Cep164* in FL cells with either MNX1 or empty vector determined by ChIP-qPCR (left panel, d). Negative control was achieved by immunoprecipitation against HA antibody, followed by detection of random region (right panel, d). Data represents mean ± SD of at least three experiments and is considered significant using two-sided student t-test (**) at $p \leq 0.01$ and (*) at $p \leq 0.05$. Western blot analysis of H3K4me1, H3K4me2, H3K4me3 and H3K27me3 in BM of leukemia mice transfected with *MNX1* ectopic expression in comparison with FL cells transfected with empty vector (Ctrl) (left panel, e), or in *invitro* FL cells (r-FL) transfected with *MNX1* ectopic expression in comparison with FL cells transfected with empty vector (Ctrl) (right panel, e). Histone H3 and Actin were used as loading control (e).

**Supplementary figure 11.** The binding of Histone H3 to MNX1 was detected by immunoprecipitation of MNX1 using HA-antibody followed by western blot using histone H3 antibody in FL cells (left panel ,a) or  BM of leukemia mice transfected with *MNX1* ectopic expression (m) in comparison to FL cells transfected with empty vector (c) (right panel, a). ETV6 with HA tag was used as a negative control. Total protein input was used as loading control (left panel, a). Most enriched pathways for differentially accessible regions of leukemia BM cells with *MNX1* ectopic expression in comparison with FL cells with empty vector using gene-ontology biological process pathways (b). In-silico predicted transcription factor binding sites by DIFFTF package from ATAC-Seq showing top differentially activated (FDR < 0.05) transcription factors (TFs) binding between MNX1 BM from leukemia mice and FL Ctrl cells (c). Scatter plot showing the correlation between differential binding regions of H3K27me3 and the differential gene expression from the RNA-Seq expression analysis (d). Results were considered at the log fold-change cutoff (logFC) of ≥ |1| and false discovery rate (FDR) of ≤0.05.

**Supplementary figure 12.** The binding of Mat2a was detected by immunoprecipitation of MNX1 using HA-antibody followed by western blot using Mat2a antibody in BM of leukemia mice transfected with *MNX1* ectopic expression (m) in comparison to FL cells transfected with empty vector (C) (a). Western blot analysis of Mat2a, Ahcy and Histone H3 in *invitro* FL cells (r-FL) transfected with *MNX1, ETV6 or MNX1::ETV6* fusion in comparison with FL cells transfected with empty vector (ctrl). Actin was used as loading control (b). Various amino acid concentration was determined by LC-ESI-MSMS from leukemia BM cells transfected with MNX1 ectopic expression normalized against total protein concentration and represented as fold difference relative to control (c). SAH concentration was determined by ELISA from *invitro* FL cells expressing MNX1 (r-FL) and leukemia BM cells (BM NSG), normalized against total protein concentration and represented as fold difference relative to control (d). Free methionine concentration was determined by LC-ESI-MSMS from FL-HSPC expressing MNX1 (MNX1-FL) and leukemia BM cells (MNX1-BM), normalized against total protein concentration and represented as fold difference relative to control (e). Recombinant Histone

H3.1& H3.3 were subjected to an invitro methyltransferase reaction using MNX1 complex pulled down with HA antibody from FL cells transfected with MNX1 (M), in comparison with FL cells with empty vector (C) in the presence of SAM and DTT. The reactions were terminated by boiling in SDS sample buffer. Separation of samples in 12% SDS-PAGE was followed by immunoblotting (IB) with H3K4me1 antibody. Reblotting was made for detection of HA and total Histone H3. As indicated, negative controls were obtained by omitting Histone H3, or pulled down immune complex (f). Quantification of the invitro methyltransferase reaction (g). Data represents mean ± SD of at least three experiments and is considered significant using two-sided student t-test (**) at $p \leq 0.01$ and (*) at $p \leq 0.05$.

**Supplement figure 13.** Data from twelve t(7;12) patients versus 62 normal human BM was selected from target cohort. Clinical attributes of the t(7;12) patients from that target cohort presenting the WBC count at diagnosis, peripheral blast percentage and cytogenetic complexity (a). Box plot showing the expression level as normalized counts from DESeq2 of the indicated genes between the t(7;12) patients and normal human BM (a). Data represents mean ± SD of at least three experiments and is considered significant using two-tailed Mann-Whitney U-test (***) at $p \leq 0.01$.

**Supplement figure 14.** Gene set enrichment analysis (GSEA) using the Gene ontology biological pathways (GO), showing GSEA enrichment plots of the common pathways between the t(7;12) patients and the MNX1 mouse model (a).

**Supplement figure 15.** ImageJ quantification of the western blot analysis of H3K4me1, H3K4me2, H3K4me3 and H3K27me3 after treatment as indicated (a). Immunofluorescence of FL cells stained with γH2AX –FITC antibody (green) and counterstained with DAPI (blue) after treatment with vehicle or 5μM Sinefungin (MNX1+S) (b). Quantification of the flowcytometry analysis of FL cells with 5μM Sinefungin (Ctrl+S, MNX+S) or without treatment (Ctrl, MNX1) after staining with DAPI for cell cycle distribution (c). Percentage of MNX1 engraftment in bone marrow of (NBSGW) in mice transplanted with FL cells, with 5μM Sinefungin (Ctrl+S, MNX+S) or without pretreatment (Ctrl, MNX1) (d). Mice were euthanized at the end and were analyzed for spleen weight, white blood cell number and hemoglobin concentration (e). mRNA expression of *MNX1* determined by quantitative real time qPCR, normalized to *Hprt* and represented as relative expression from FL cells with 5μM Sinefungin (Ctrl+S, MNX+S) or without treatment (Ctrl, MNX1) (f). Data represents mean ± SD of at least three experiments and is considered significant using two-sided student t-test between MNX1 and MNX1+S (**) at $p \leq 0.01$ and (*) at $p \leq 0.05$.

**Supplement figure 16.** Percentage of GFP/YFP chimerism in peripheral blood of (NBSGW) mice transplanted with FL with 5μM Sinefungin (Ctrl+S, MNX+S) or without treatment (Ctrl, MNX1), as determined by flowcytometry every two weeks (a). Data represents mean ± SD of at least three experiments and is considered significant using two-sided student t-test between MNX1 and MNX1+S (**) at $p \leq 0.01$ and (*) at $p \leq 0.05$. LogFC heat map of downregulated (blue) and upregulated (red) differentially expressed genes of MNX1 that are altered by Sinefungin treatment (b). Differentially expressed genes were selected based on a LogFC1.5 and FDR ≤ 0.05. Schematic diagram for the proposed model of action of MNX1 (c).

GFP: Green fluorescence protein, YFP: Yellow fluorescence protein

a

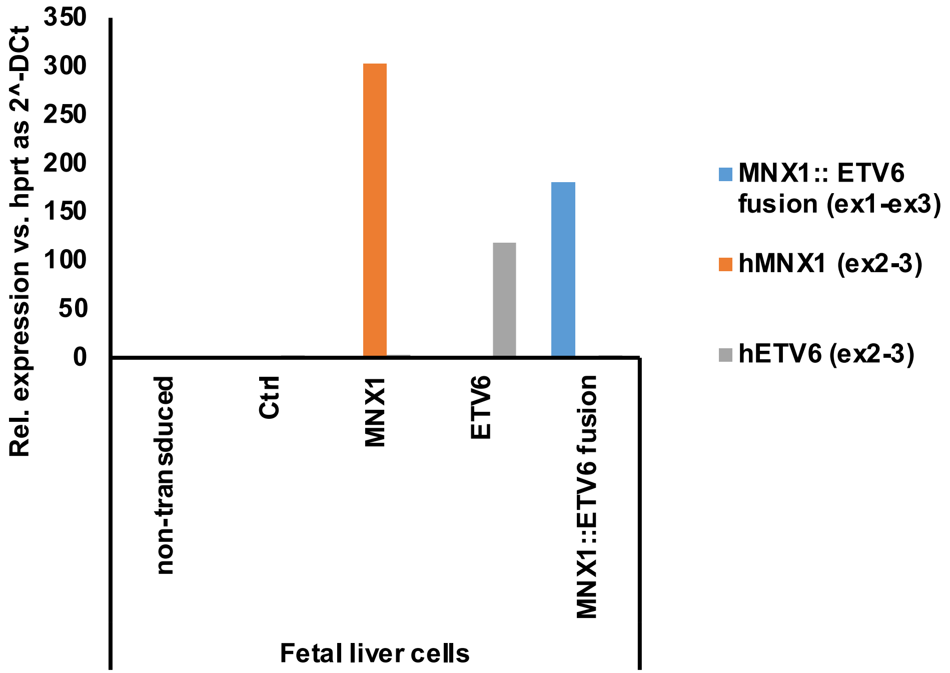


b

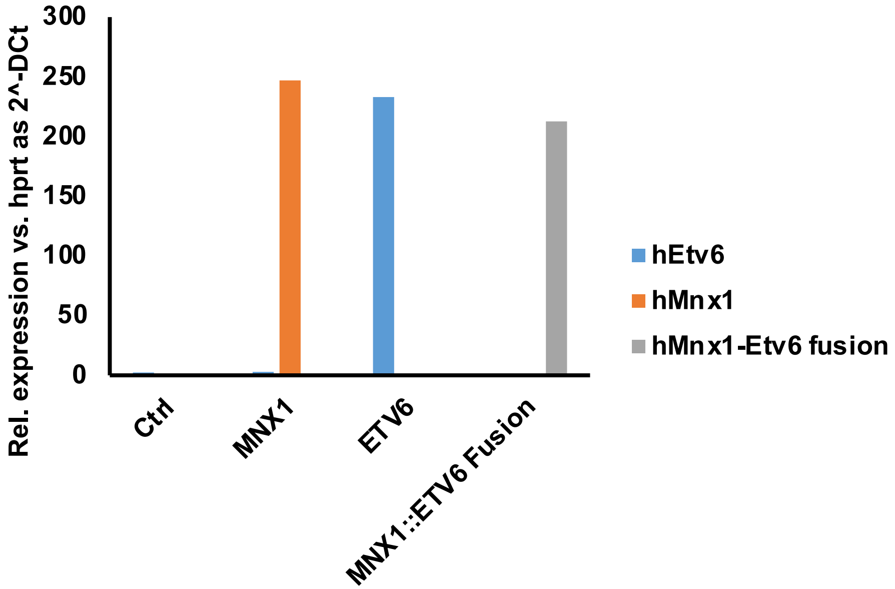


c

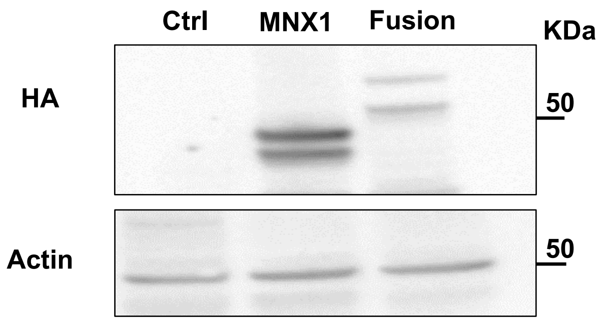


Supplementary Figure 1

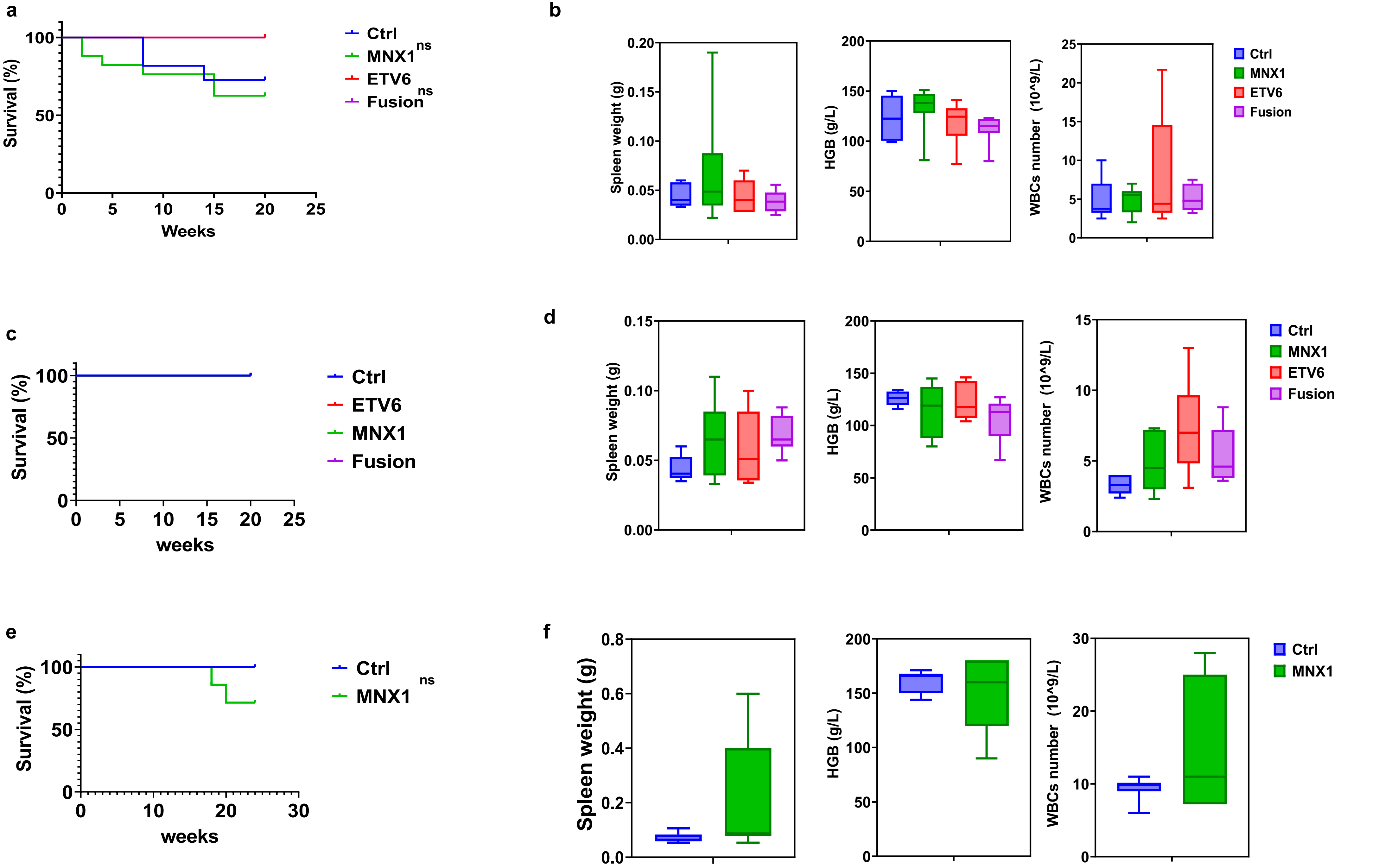


Supplementary Figure 2

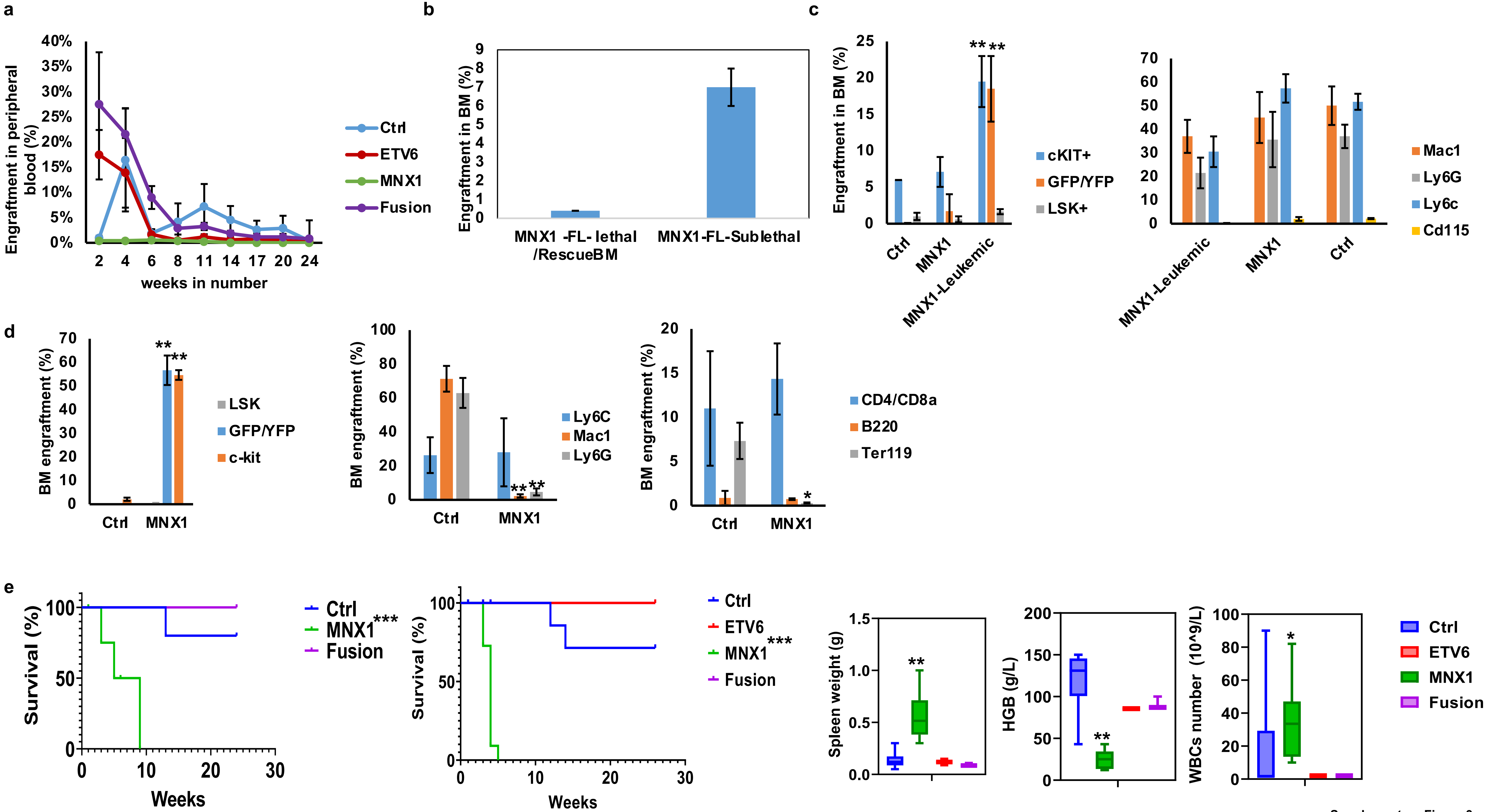


Supplementary Figure 3

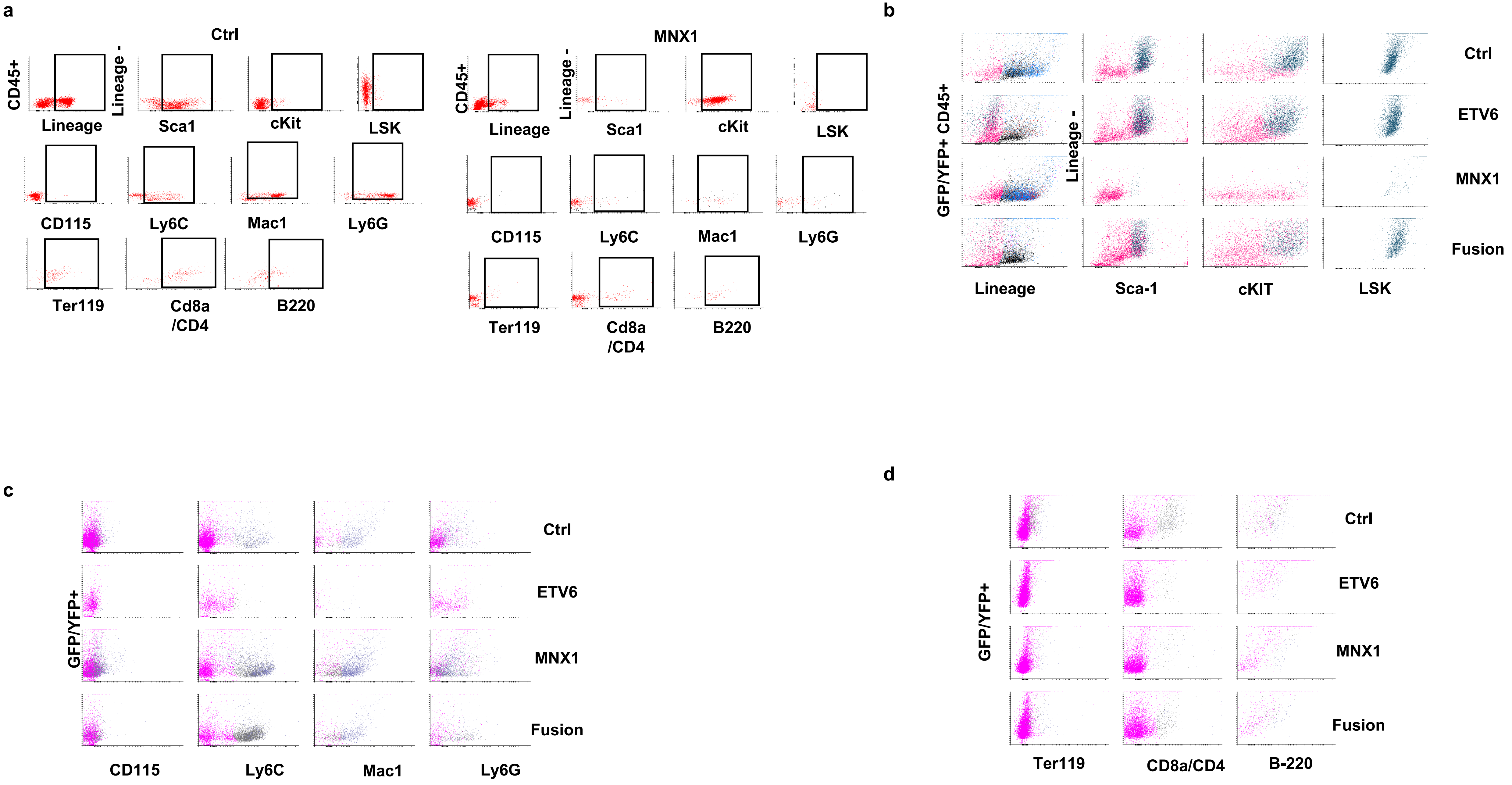


Supplementary Figure 4

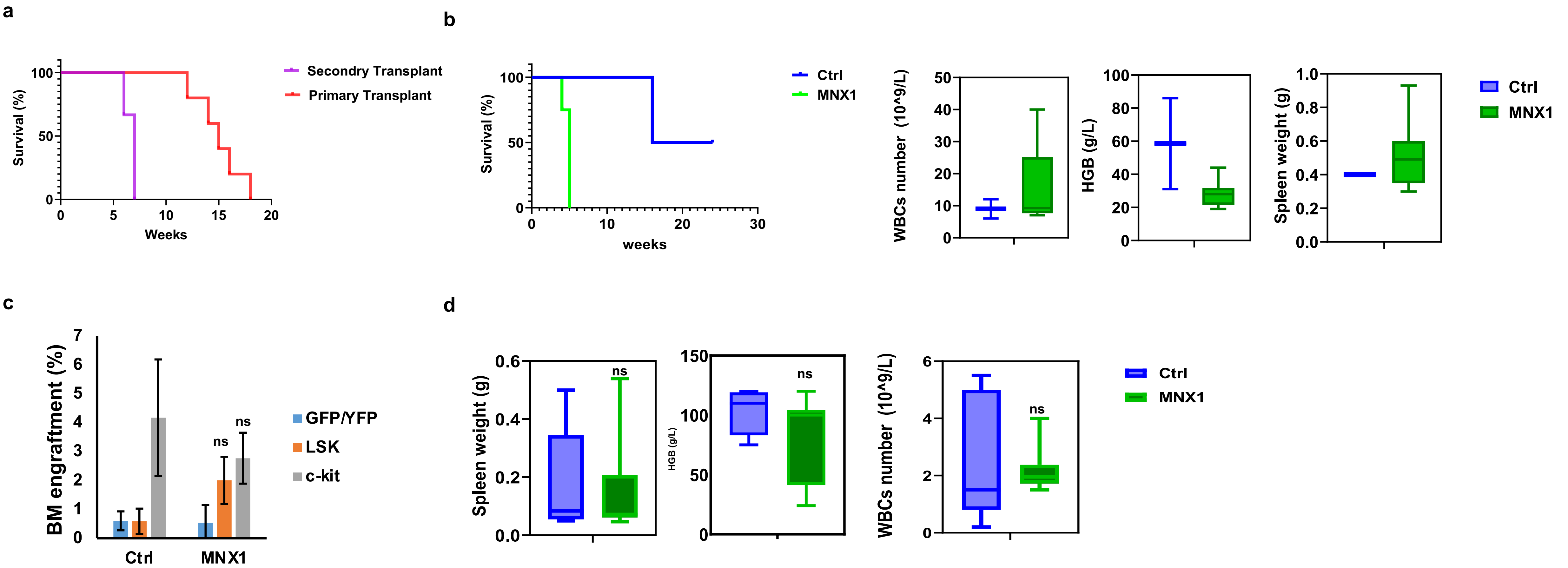


Supplementary Figure 5

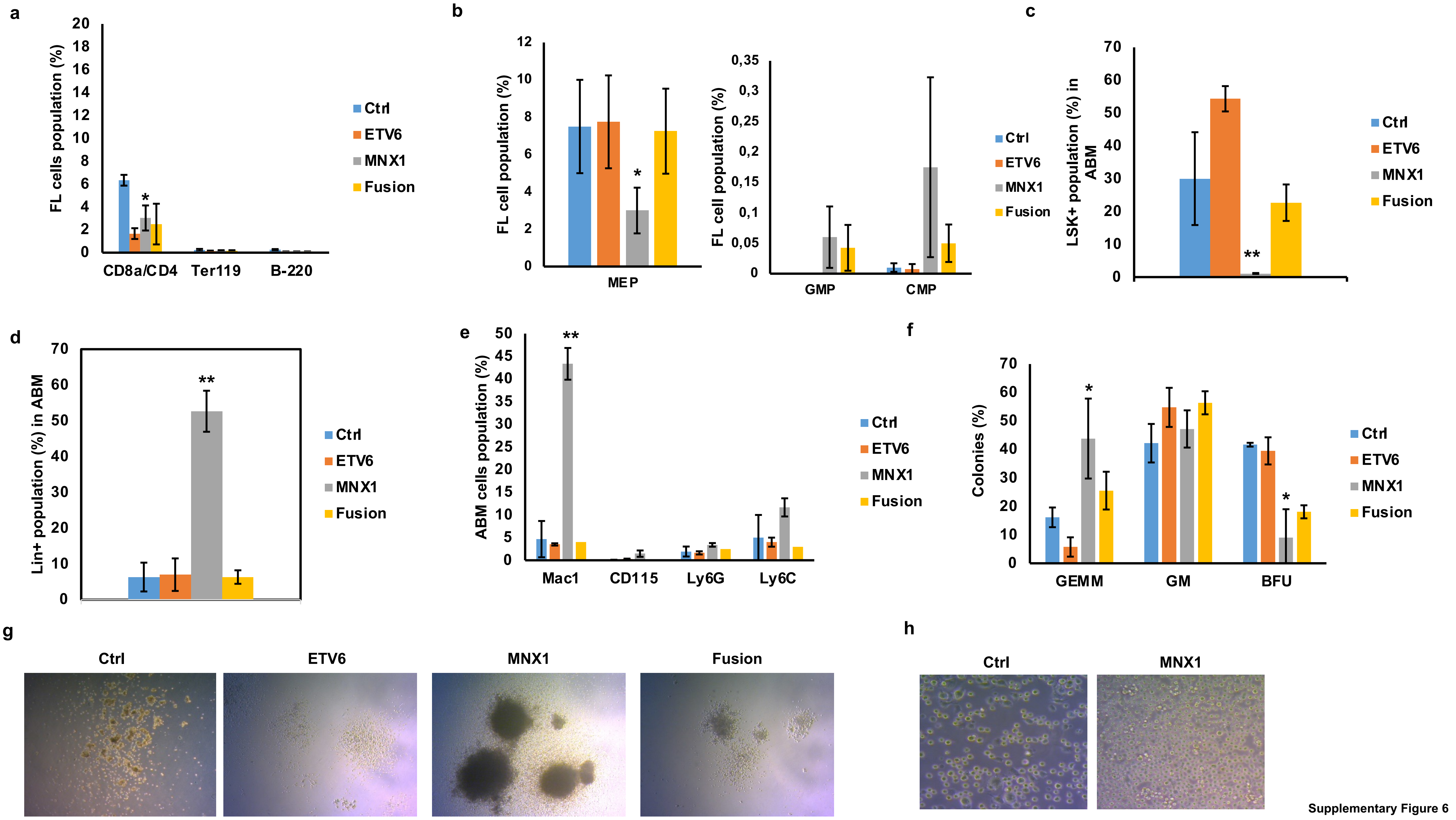


Supplementary Figure 6

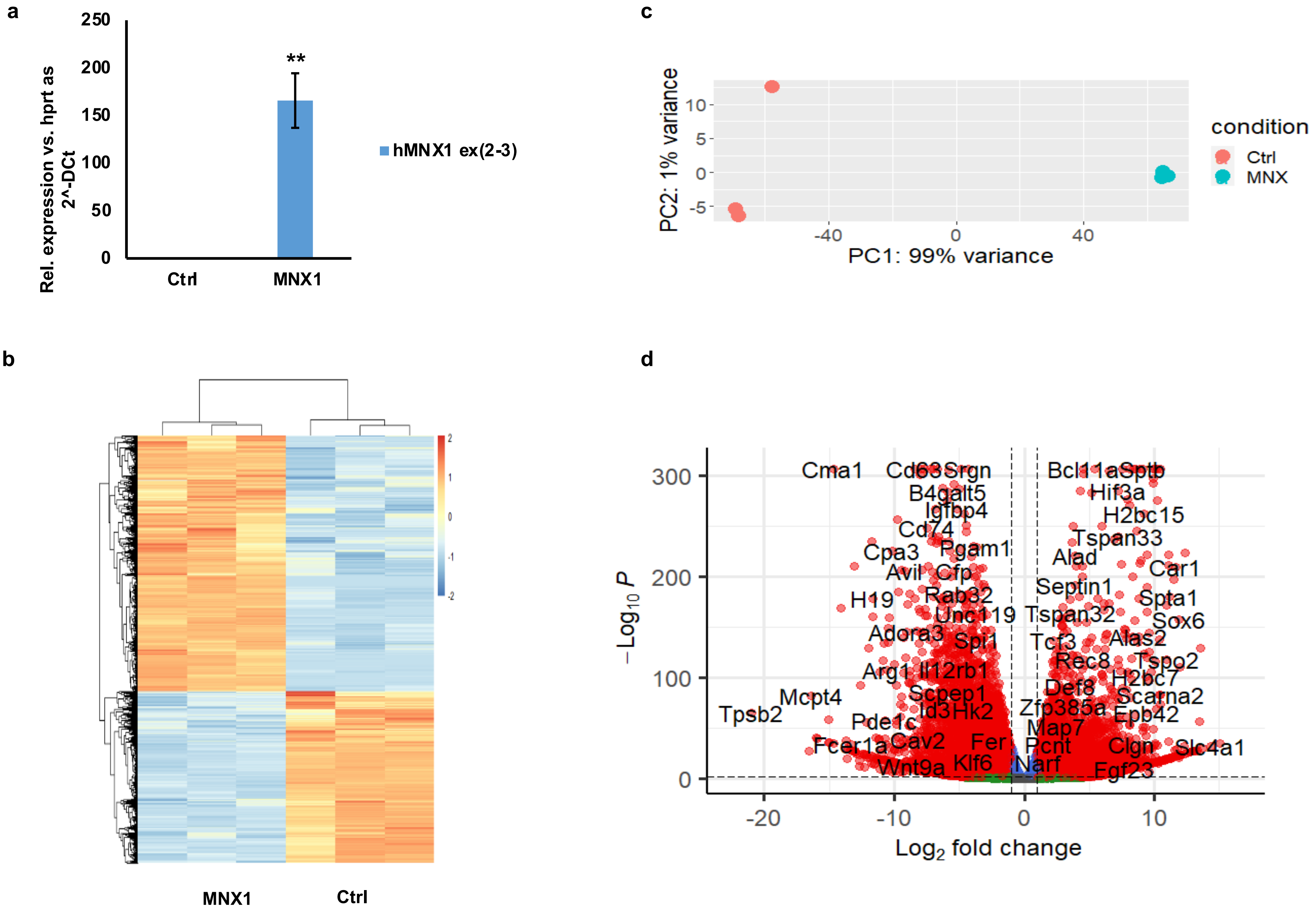


Supplementary Figure 7

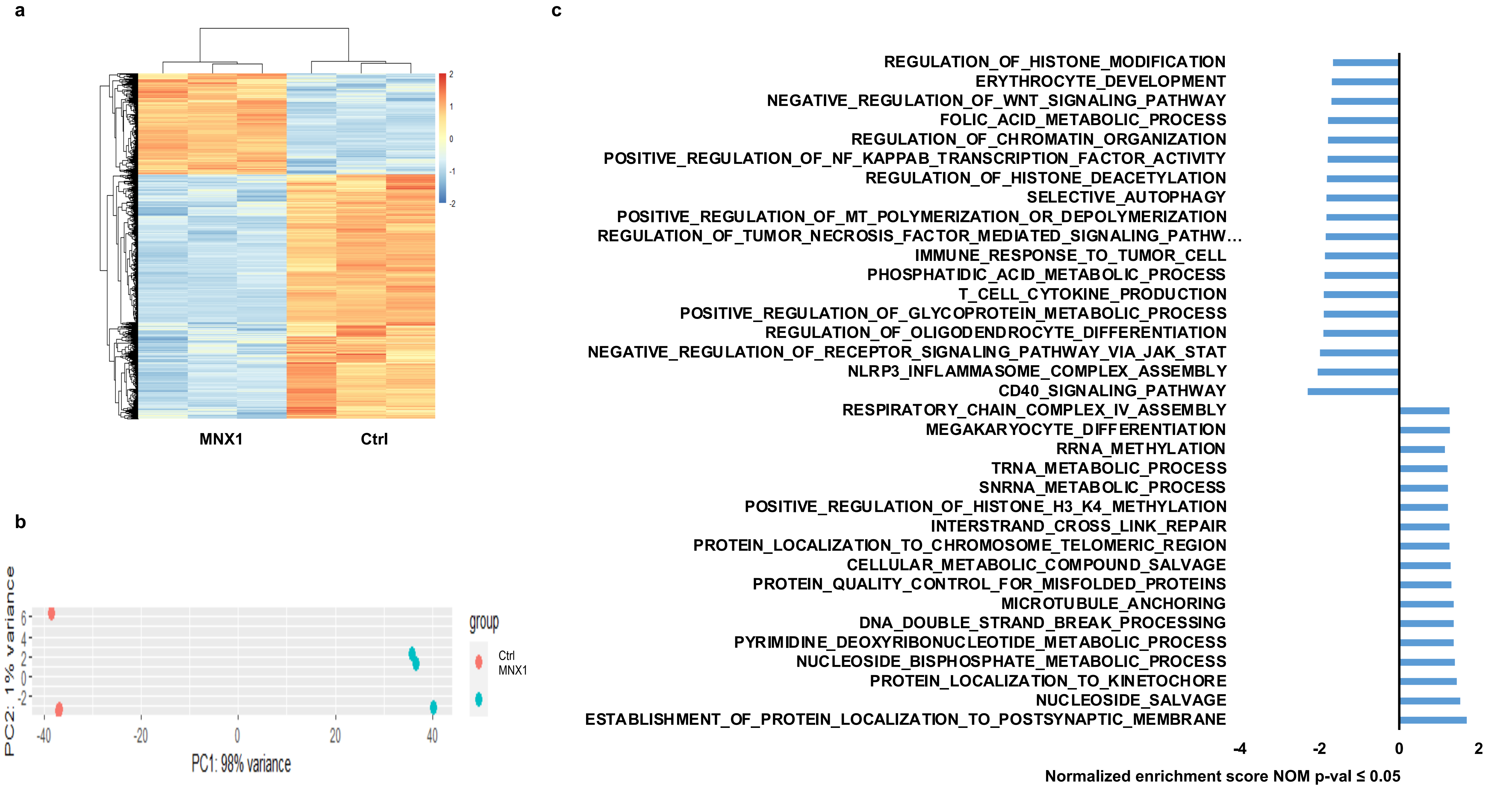


Supplementary Figure 8

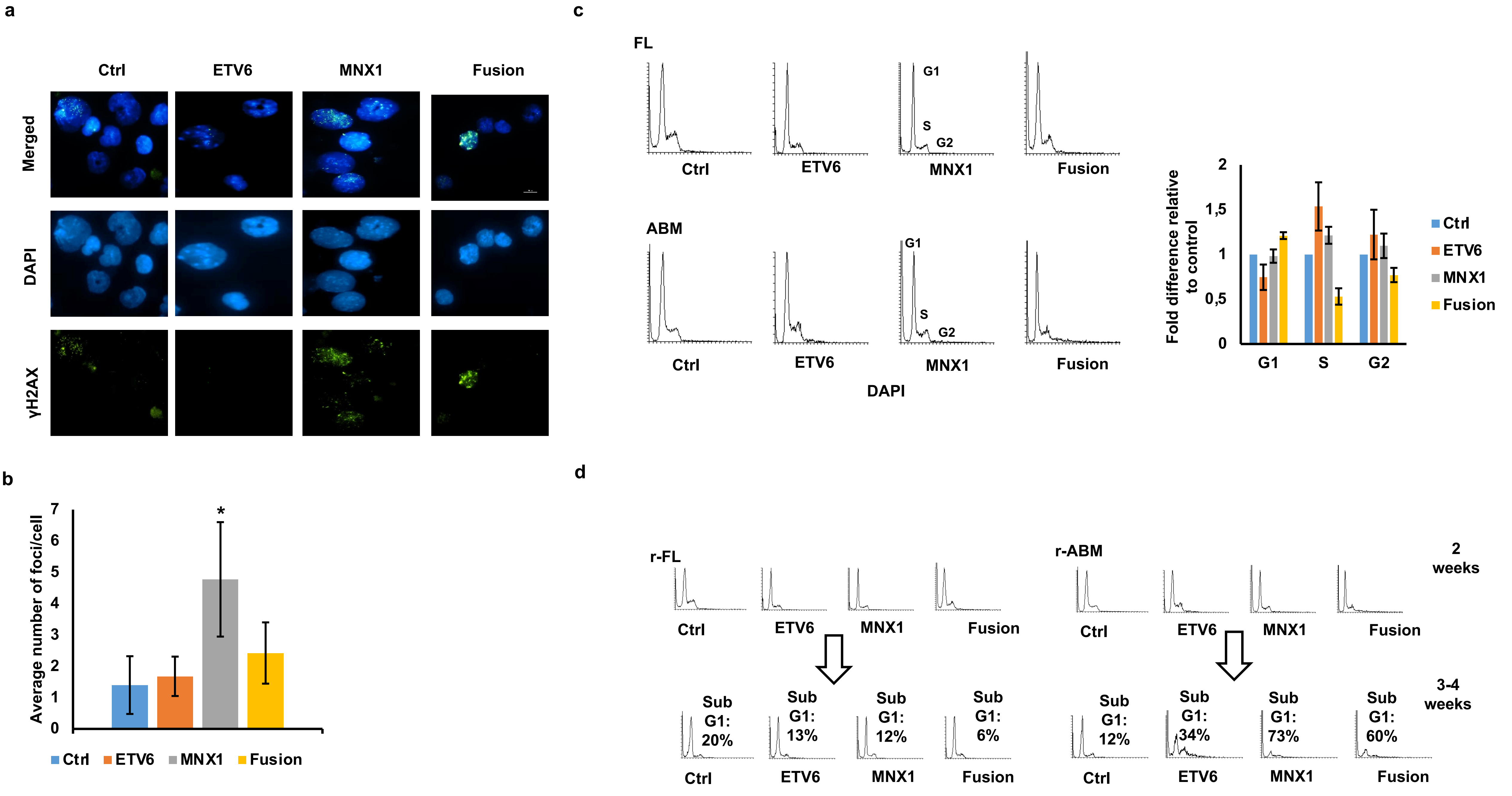


Supplementary Figure 9

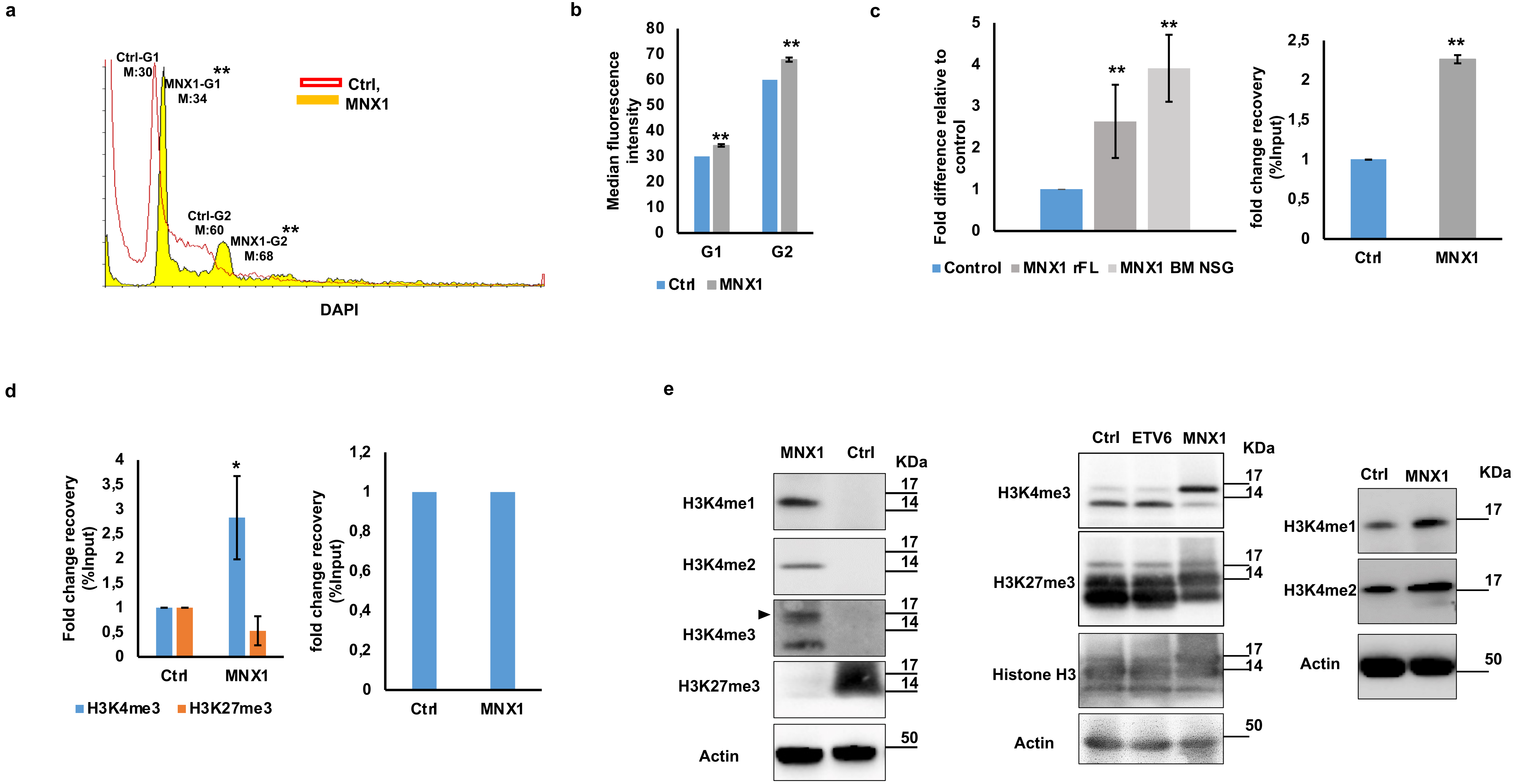


Supplementary Figure 10

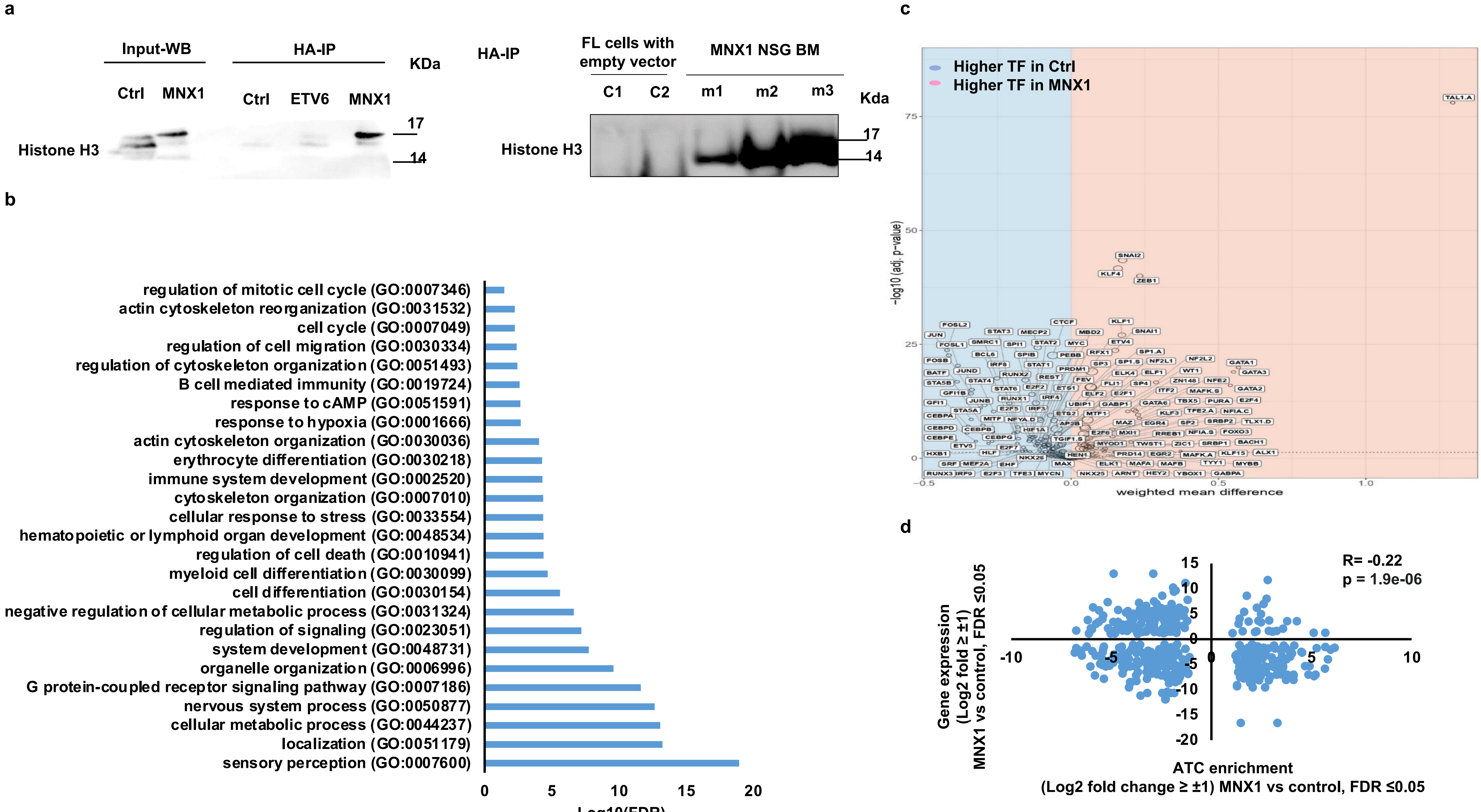


Supplementary Figure 11

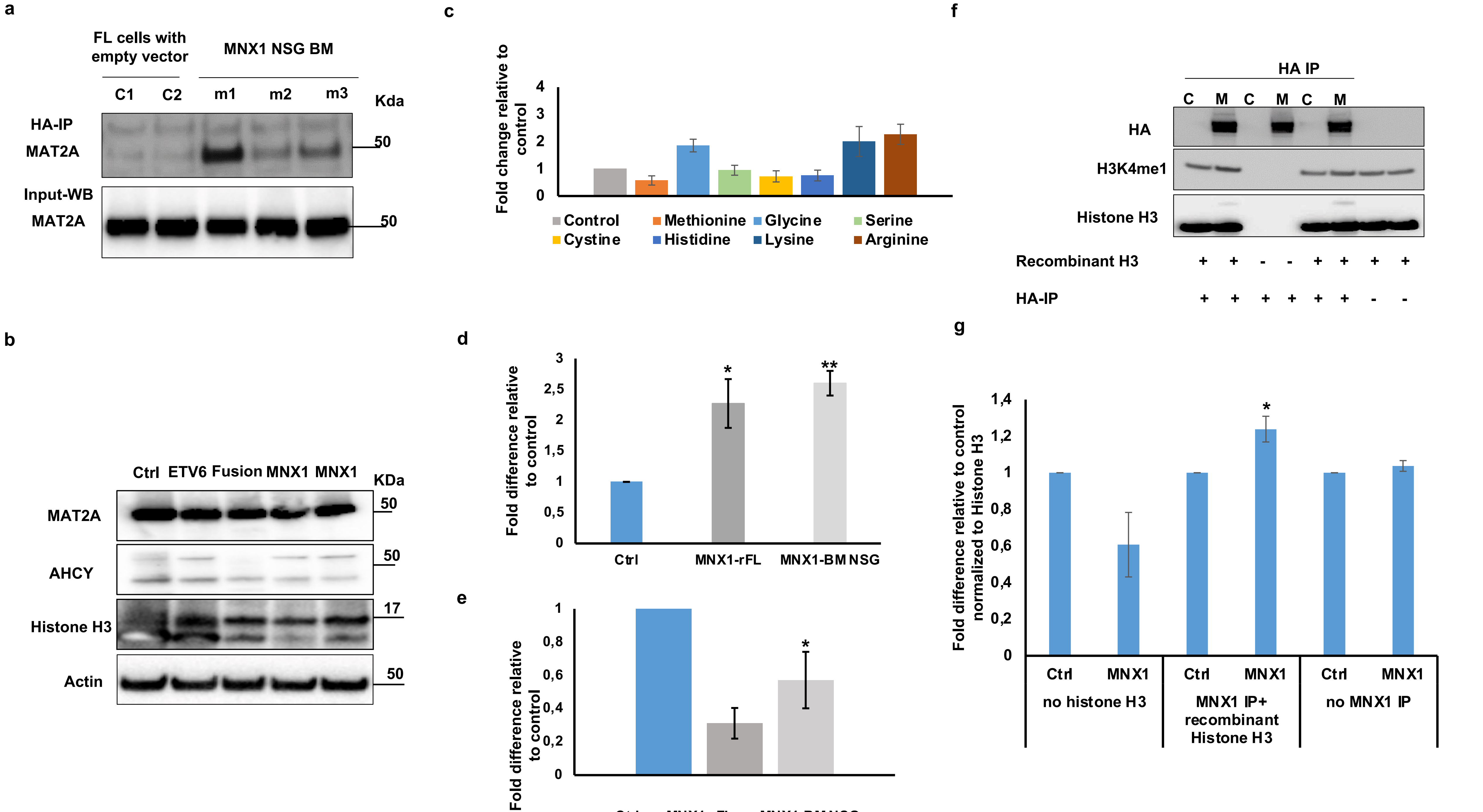


Supplementary Figure 12

a

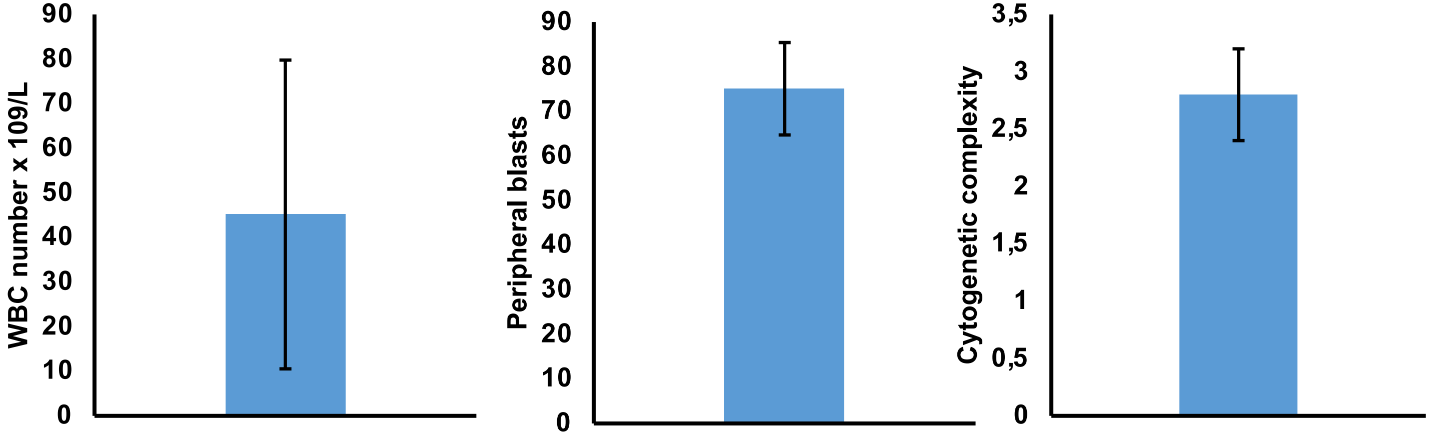

WBC number x 109/L
90
80
70
60
50
40
30
20
10
0
Peripheral blasts
90
80
70
60
50
40
30
20
10
0
Cytogenetic complexity
3,5
3
2,5
2
1,5
1
0,5
0


b

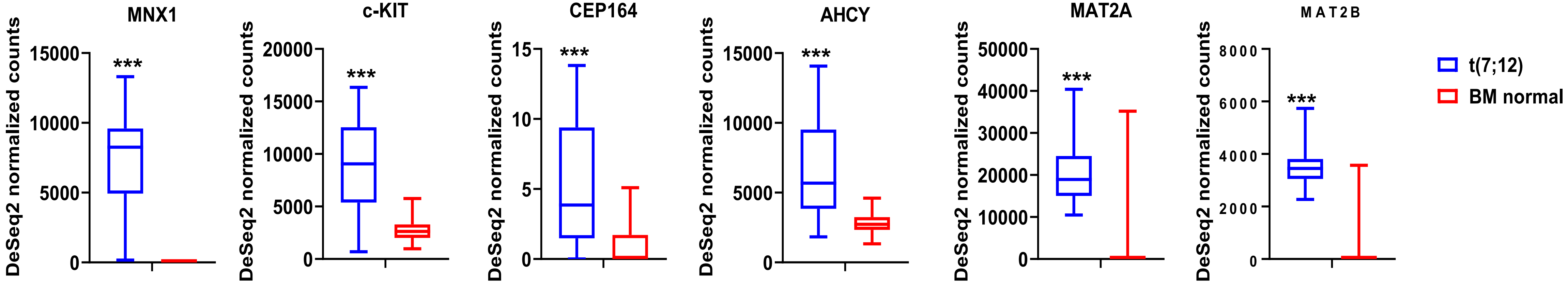

MNX1
c-KIT
CEP164
AHCY
MAT2A
MAT2B
DeSeq2 normalized counts
15000
10000
5000
0
20000
15000
10000
5000
0
15
10
5
0
15000
10000
5000
0
50000
40000
30000
20000
10000
0
8000
6000
4000
2000
0
***
t(7;12)
BM normal

a

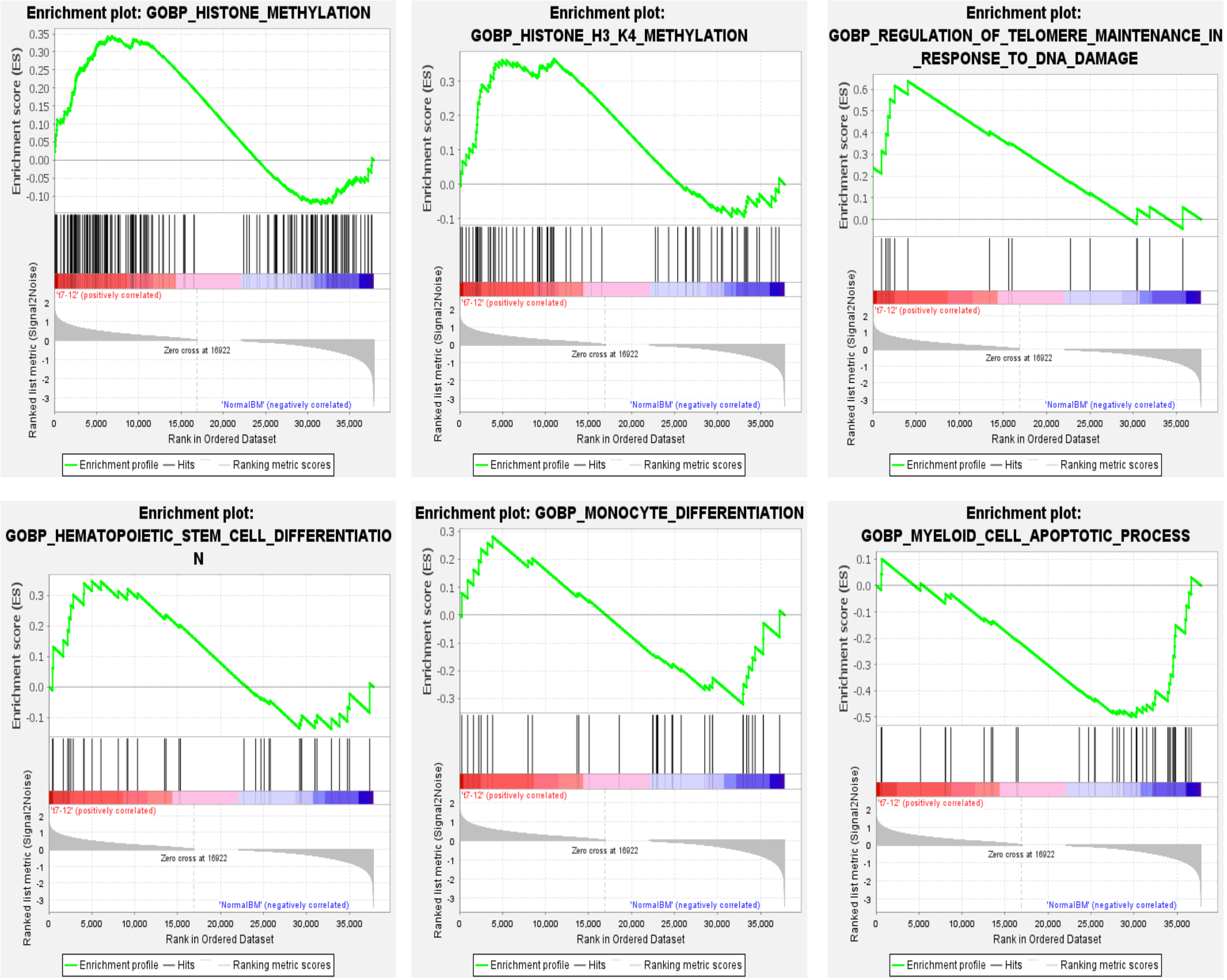


Supplementary Figure 14

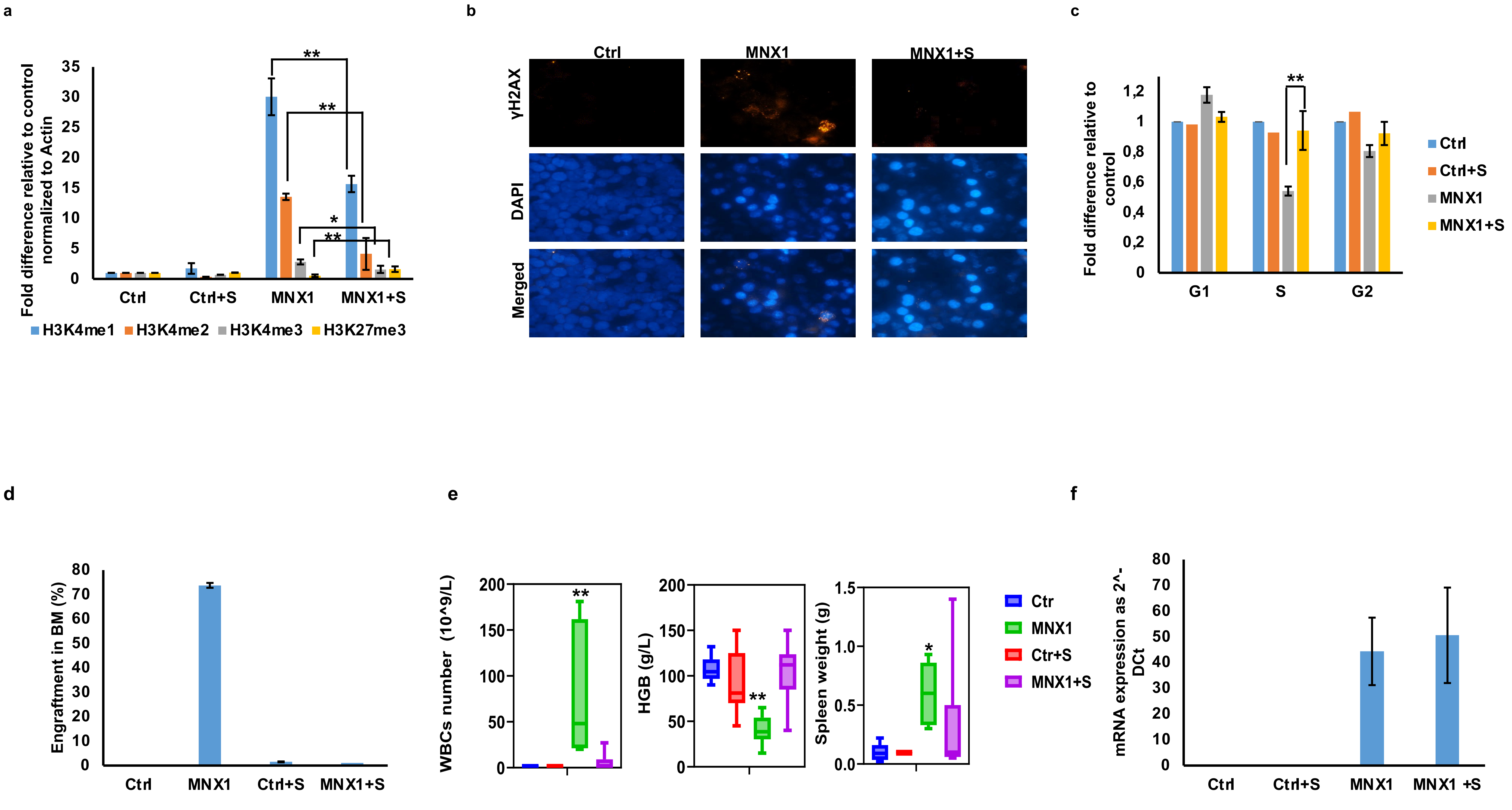


Supplementary Figure 15

a

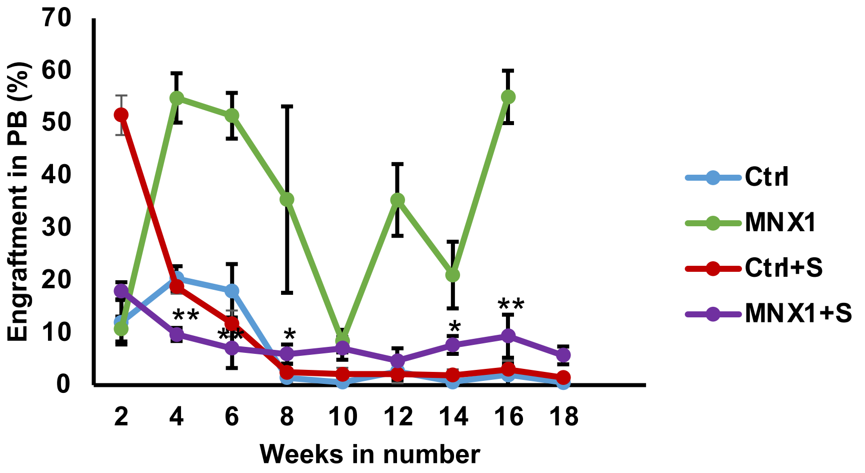


b

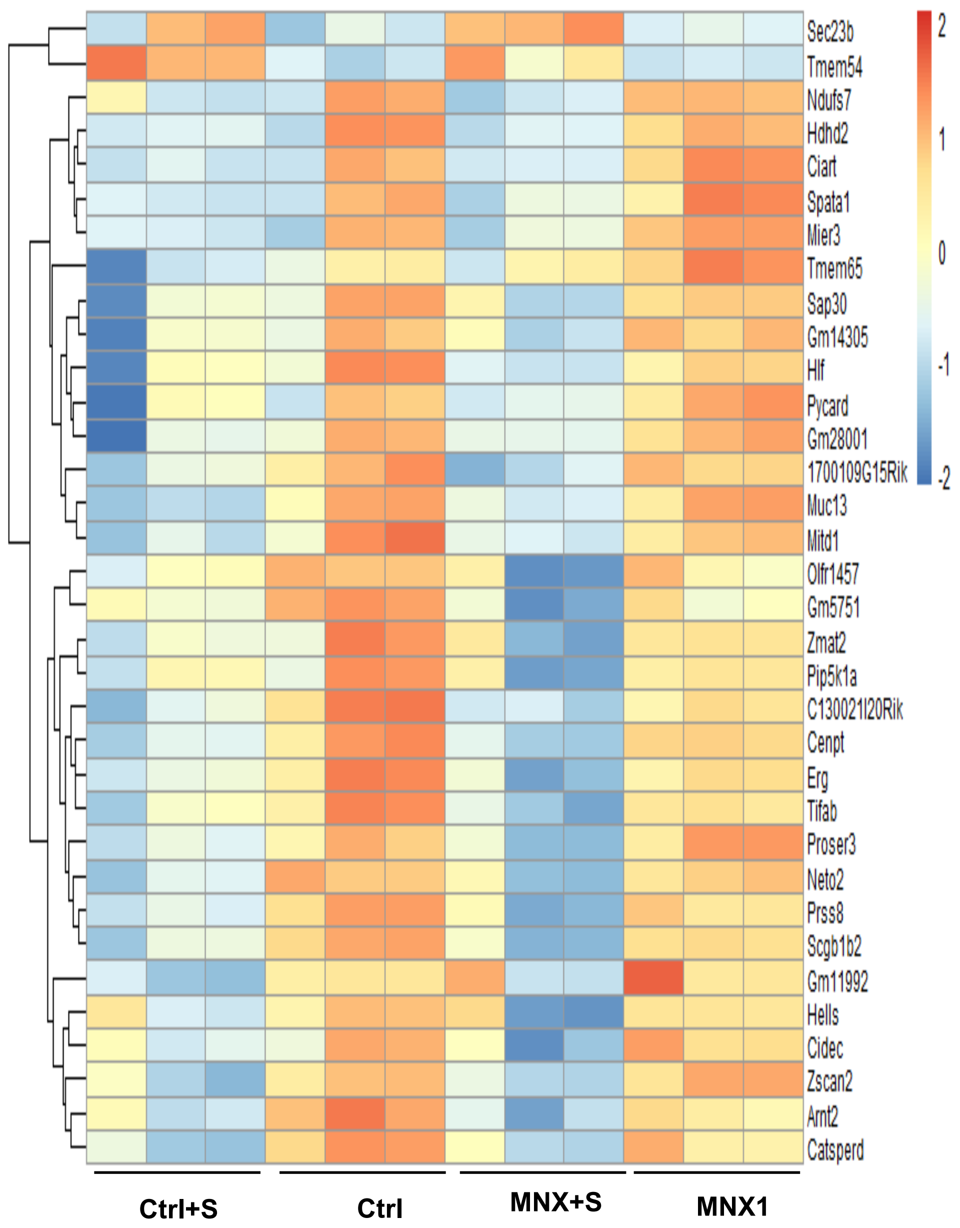


c

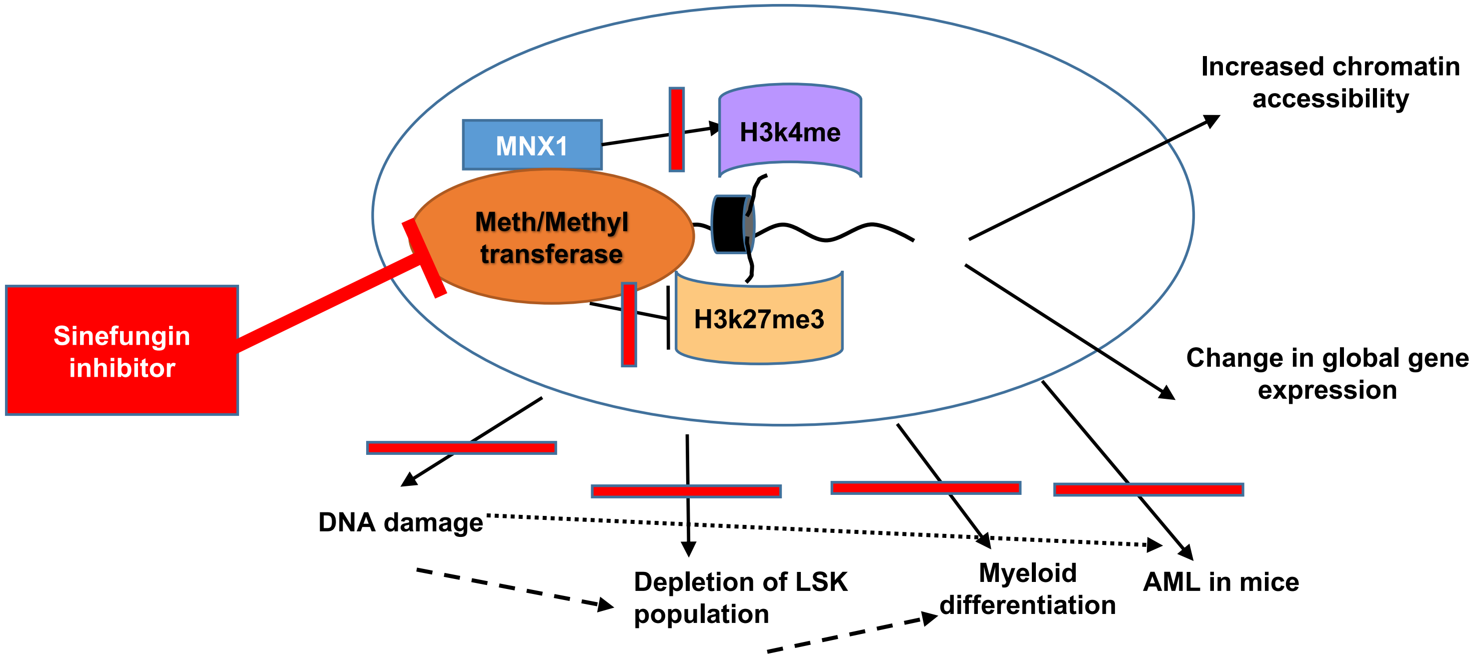


Supplementary Figure 16
